# Lactylation of Influenza Virus Polymerase Acidic Protein Promotes Viral Replication and Pathogenicity

**DOI:** 10.64898/2026.07.10.737663

**Authors:** Shaoyu Tu, Yue Du, Wei Liang, Xiaoyi Xu, Jiahui Zou, Yuchao Yang, Chuhan Xiong, Yanglin Li, Meijun Jiang, Aotian Ouyang, Tong Chen, Meilin Jin, Huanchun Chen, Hongbo Zhou

## Abstract

Influenza virus poses a potential risk of triggering the next global pandemic. In-depth investigation into the mechanisms underlying influenza virus replication and pathogenicity will provide robust support for controlling influenza virus infection. Although post-translational modifications are known to regulate viral infection, the role of lactylation in influenza virus replication remains elusive. In this study, influenza virus ribonucleoprotein complex subunits are found to be lactylated. Specifically, ATAT1 promotes viral polymerase acidic protein (PA) lactylation and enhances viral replication. In contrast, SIRT1 mediates de-lactylation of PA and exerts an inhibitory effect on viral replication. Further investigations reveal lactylation of PA at residues K605 and K609 is essential for viral replication and pathogenicity. Mechanistically, PA K605/609 residues are localized at the interaction interface of the ANP32-mediated polymerase asymmetric dimer; mutation at these residues inhibits polymerase asymmetric dimerization, thereby impairing RNA production during viral genome replication. Collectively, this study uncovers a novel mechanism by which influenza virus hijacks host enzymes to mediate PA lactylation, and expands the molecular regulatory network of influenza virus infection.

## Introduction

Influenza virus is a segmented, negative-sense, single-stranded RNA virus classified into types A, B, C, and D based on the antigenicity of its nucleoprotein (NP) and matrix protein 1 (M1) (1). Of these, influenza A virus poses the greatest threat to public health, causing 290,000-650,000 human deaths annually. The viral core is composed of eight viral ribonucleoprotein complexes (vRNPs), the minimal replication unit of the virus, whose functions dictate viral replication efficiency and pathogenicity (2). During the early phase of infection, the viral polymerase complex exerts transcriptional activity to generate mRNA for *de novo* viral protein synthesis (3). With the progression of infection, newly synthesized vRNP subunits translocate into the nucleus to initiate genome replication, which proceeds in two sequential steps. Initially, cRNA intermediates are synthesized using parental vRNAs as templates, followed by the generation of progeny vRNAs from cRNA templates (4). During this process, viral polymerase complex undergoes dimerization and adopts multiple conformational states, primarily asymmetric and symmetric dimers. The ANP32-bridged asymmetric dimer is critical for both cRNA and vRNA synthesis, whereas the symmetric dimer is responsible for vRNA production (5–7).

Influenza virus encodes only a limited set of proteins and thus employs multiple strategies to diversify its protein functions, a key one of which is the regulation of post-translational modifications (PTMs) (8). Such modifications enable the virus to dynamically modulate protein functions without *de novo* protein synthesis. Accumulating studies have demonstrated that influenza virus hijacks various PTMs, including glycosylation, acetylation, ubiquitination, phosphorylation, and SUMOylation, to facilitate multiple stages of its replication cycle (9–14). Specifically, multiple glycosylation sites have been identified within the viral hemagglutinin (HA), which is critical for regulating HA receptor specificity and affinity (15, 16). During uncoating, influenza virus exploits the host E3 ligase Itch to mediate ubiquitination of viral M1, leading to proteasomal degradation of M1 and promoting cytoplasmic release of vRNPs (17). At late stage of infection, viral non-structural protein 2 (NS2) blocks AIMP2-mediated M1 ubiquitination while enhancing M1 SUMOylation at residue 242; this drives the assembly of the vRNP-M1-NS2 complex and promotes vRNP nuclear export (18). Despite these findings, the complete repertoire of PTMs on influenza viral proteins eludes full characterization.

With advances in detection technologies, emerging PTMs have been successively identified. Histone lactylation was reported to regulate the expression of homeostasis-related genes in 2019, and subsequent studies have demonstrated the widespread prevalence of lactylation on non-histone proteins (19–21). Recent evidence indicates that certain viruses disrupt host glycolysis and hijack the lactylation machinery to facilitate viral replication. For instance, porcine reproductive and respiratory syndrome virus (PRRSV) infection induces lactate accumulation, which upregulates histone H3K18 lactylation to promote HSPA6 expression and ultimately suppresses host innate immunity (22). White spot syndrome virus (WSSV) enhances lactylation of host histones at H3K18 and H4K12 to upregulate ribosomal protein S6 kinase 2 (S6K2) expression to promote efficient viral replication (23). Beyond histone lactylation, Kaposi’s sarcoma-associated herpesvirus (KSHV) infection promotes NAT10 lactylation, thereby enhancing NAT10-mediated N4-acetylcytidine (ac4C) modification of tRNA(ˢᵉʳ)-CGA-1-1, which increases translational efficiency of viral mRNAs and drives progeny virion production and viral reactivation (24). Most recently, lactylation of the SARS-CoV-2 spike (S) protein was shown to enhance its binding to the ACE2 receptor, expanding our understanding of how viruses exploit host PTM systems to augment the functional versatility of their own proteins (25). However, whether influenza virus orchestrates its replication through lactylation of host or viral proteins remains poorly explored.

Transcriptomic and metabolomic analyses have established that influenza virus infection upregulates cellular glycolysis and leads to elevated lactate levels (26–28). While lactate is known to suppress host innate immunity by blocking MAVS aggregation and subsequent signal transduction (29, 30), its direct effects on influenza virus replication require further investigation. Notably, treatment with the glycolysis inhibitor 2-deoxy-D-glucose (2-DG) prolongs viral mRNA synthesis and impairs genomic vRNA accumulation (31, 32). Nevertheless, the precise mechanism by which glycolysis and its metabolic products regulate influenza viral polymerase function has yet to be fully elucidated.

In this study, we demonstrate that lactate promotes influenza virus replication. Further investigations reveal that subunits of viral vRNP complex undergo lactylation. ATAT1 promotes, whereas SIRT1 inhibits lactylation of viral PA protein, thereby exerting positive and negative regulatory effects on influenza virus replication, respectively. Multiple lactylated lysine residues were identified on the viral polymerase acidic protein, with residues K605 and K609 being markedly crucial for viral replication and pathogenicity. Lactylation at these two residues facilitates the formation of ANP32-mediated polymerase asymmetric dimer, thus promoting RNA production during viral genomic replication. These findings not only expand the molecular regulatory network underlying influenza virus infection, but also provide promising targets for the development of host-directed preventive and therapeutic strategies against influenza.

## Materials and methods

### Cells and Viruses

Human lung adenocarcinoma A549 cells and Madin-Darby canine kidney (MDCK) cells were cultured in Dulbecco’s modified Eagle’s medium (DMEM), human embryonic kidney 293T (HEK293T) cells were maintained in Roswell Park Memorial Institute 1640 (RPMI 1640) medium. All cell lines were incubated at 37 ℃ in a humidified 5% CO₂ atmosphere, with their respective culture media supplemented with 10% fetal bovine serum (FBS) and 1% penicillin-streptomycin.

Wild-type and PA-K605A/K605R/K609A mutant influenza A virus (A/Puerto Rico/8/34; PR8, H1N1) were rescued using an eight-plasmid reverse genetics system, following a previously described protocol (33). Viral titers were quantified on MDCK cells, and the resulting viral stocks were aliquoted and stored at -80 ℃ for subsequent use.

### Antibodies and Reagents

Primary antibodies used in this study included the rabbit Pan-L-lactyl lysine (Pan-Kla) antibody (PTM-1401RM, PTM BioLab, China), rabbit anti-LDHA (19987-1-AP, Proteintech, USA), mouse anti-β-actin (66009-1-Ig, Proteintech, USA), mouse anti-GAPDH (60004-1-Ig, Proteintech, USA), rabbit anti-SIRT1 (13161-1-AP, Proteintech, USA), rabbit anti-ATAT1 (28828-1-AP, Proteintech, USA), rabbit anti-influenza A virus PA protein (GTX644074, GeneTex, USA), rabbit anti-influenza A virus PB1 protein (GTX125923, GeneTex, USA), rabbit anti-influenza A virus PB2 protein (GTX125926, GeneTex, USA), rabbit anti-influenza A virus NP protein (GTX636247, GeneTex, USA), mouse monoclonal antibodies against HA tag (66006-2-Ig, Proteintech, USA) and FLAG tag (66008-4-Ig, Proteintech, USA), as well as mouse anti-ANP32A (67687-1-Ig, Proteintech, USA) and anti-ANP32B (66160-1-Ig, Proteintech, USA).

Secondary antibodies comprised horseradish peroxidase (HRP)-conjugated goat anti-mouse IgG (AS003, Abclonal, USA), HRP-conjugated goat anti-rabbit IgG (AS014, Abclonal, USA), Cy3-conjugated goat anti-rabbit IgG (AS007, Abclonal, USA) and FITC-conjugated goat anti-mouse IgG (AS001, Abclonal, USA).

The chemical reagents employed were as follows: Oxamate (HY-W013032, MedChemExpress, MCE, USA), Galloflavin (HY-W040118, MCE, USA), lactic acid (HY-B2227, MCE, USA), sodium lactate (HY-B2227B, MCE, USA), N-p-Tosyl-L-phenylalanine chloromethyl ketone (TPCK)-treated trypsin (232-650-8, Sigma-Aldrich, USA), Lipofectamine 2000 transfection reagent (11668019, Thermo Fisher Scientific, USA) and 4′,6-diamidino-2-phenylindole (DAPI) staining solution (C1002, Beyotime, China).

### Plasmids and Transfection

Genes encoding ESCO1 (NM_052911.3), ESCO2 (NM_001017420.3), ATAT1 (NM_001413067.1), MYST1 (NM_032188.3), AARS1 (NM_001605.3), SIRT1 (NM_012238.5), SIRT2 (NM_012237.4), SIRT5 (NM_012241.5), SIRT6 (NM_016539.4), SIRT7 (NM_016538.3), HDAC1 (NM_004964.3), HDAC2 (NM_001527.4) and HDAC3 (NM_003883.4) were cloned into the pCAGGS-HA vector using restriction sites Cla I/Kpn I (ESCO1, ATAT1, MYST1, AARS1, HDAC1, HDAC2, SIRT6, SIRT7), EcoR I/Xho I (SIRT5, HDAC3), or Cla I/Nhe I (ESCO2, SIRT1). Eukaryotic expression plasmids for viral proteins (PA, PB1, PB2, NP in pcDNA3.1 vector, FLAG/HA-tagged PA, FLAG/HA-tagged PB1) and plasmids for viral minigenome assays (Renilla luciferase control plasmid pRL-TK, viral reporter plasmid pPol I-luc) were preserved in our laboratory. Site-directed mutagenesis was performed to construct the mutant plasmids: PA carrying alanine- or arginine-substituted mutations at residues 356, 358, 361, 362, 605, or 609, PA-C95A, PA-D108A and ATAT1-D157N. The HA-tagged SIRT1 catalytic domain deletion mutant (SIRT1-ΔCata) plasmid was generated by homologous recombination. Pol I plasmids expressing influenza virus segment 5 vRNA or cRNA were constructed using a pHW2000 vector with the CMV and T7 promoters deleted. Plasmids for split-luciferase assays (PB1-Gluc-N, PB1-Gluc-C, ANP32A-Gluc-C, ANP32B-Gluc-C) were constructed as detailed previously. All plasmids constructed in the present study were validated via Sanger sequencing.

A549 and HEK293T cells were transfected with indicated siRNAs or plasmids using Lipofectamine 2000 transfection reagent in accordance with the manufacturer’s recommended protocol. Transfected cells were further cultured for 18-24 h before subsequent experiments. The primers for plasmid construction and the LDHA-, ATAT1- and SIRT1-targeting siRNAs are detailed in Table S2.

### Virus infection and titration

Influenza virus infection was conducted as described previously (34). Briefly, A549 cells in 12-well plates were rinsed twice with PBS and then incubated with influenza virus at the indicated MOI for 1 h at 37 ℃. Following viral adsorption, the cells were re-rinsed twice with PBS and cultured in DMEM supplemented with 0.25 µg/mL TPCK-treated trypsin at 37 ℃ under standard incubation conditions. Supernatants were harvested at designated time points post-infection. For viral titration, MDCK cells were inoculated with 10-fold serially diluted virus. After absorption, cells were cultured in medium containing 0.25 µg/mL TPCK-treated trypsin for 72 h. Viral titers were determined by TCID₅₀ assays and calculated using the Reed-Muench method.

### Cell viability assay

Cell viability was measured using a CCK-8 assay kit (GK10001, GlpBio; USA). A549 cells were seeded in 96-well plates, and treated with various concentrations of test compounds for 24 h after reaching 80%-90% confluence (n = 8 per group). Then 10 μL of CCK-8 solution was added to each well, followed by incubation at 37 °C for 1 h in the dark. Absorbance at 450 nm was recorded using a TECAN SPARK microplate reader.

### Minigenome Assay

HEK293T cells in 12-well plates were transfected with 0.15 μg wild-type or mutant PA, 0.3 μg PB1, 0.3 μg PB2, 0.6 μg NP (all in pcDNA3.1 vector), 0.3 μg viral reporter plasmid pPol I-luc and 20 ng pRL-TK (Renilla luciferase) when cell confluency reached 70-80%. At 24 h post-transfection, cells were lysed with 250 μL passive lysis buffer (Promega) for 30 min on ice. Firefly and Renilla luciferase activities were assayed with the Dual-Luciferase Reporter Assay System (E1910, Promega, USA) on a TECAN SPARK microplate reader, in accordance with the manufacturer’s protocol. Relative viral polymerase activity was normalized to Renilla luciferase activity as internal control. For the viral polymerase activity complementation assay, HEK293T cells were transfected with 0.075 μg mutant PA and complemented with 0.075 μg PA-C95A or PA-D108A; all other plasmids were co-transfected as described above.

### Quantitative Reverse Transcription-PCR (qRT-PCR)

To assess the effect of PA mutations on viral RNA synthesis, A549 cells in 12-well plates were inoculated with WT or PR8 mutants carrying PA K605A, PA K605R, or PA K609A substitutions at a MOI of 1. Cells were collected at 2, 5 and 8 hours post-infection. The levels of viral segment 5 mRNA, cRNA, and vRNA were quantified as previously reported (35). Briefly, total cellular RNA was extracted using TRIzol reagent. Tag-specific primers for viral segment 5 mRNA, cRNA or vRNA, or an unmodified oligo(dT) primer for the internal reference gene GAPDH were used for reverse transcription with a commercial RT kit (6210A, Takara, Japan). The resulting cDNAs were 10-fold serially diluted, and quantitative PCR (qPCR) was performed on a ViiA7 Real-Time PCR System (ABI, USA) using 2× Universal SYBR Green Fast qPCR Mix (RK21203; ABclonal Technology, USA). Relative viral RNA levels were normalized to internal control GAPDH and calculated using the 2⁻ΔΔCT method. Fold changes in viral RNA levels were further calibrated to the corresponding RNA levels at 2 h post-infection for each experimental group.

To assess the effect of PA mutations on viral RNA synthesis under transfection conditions, HEK293T cells in 12-well plates were co-transfected with 0.15 μg of WT PA or PA variants containing K605A, K605R, or K609A substitutions, 0.3 μg each of PB1 and PB2, 0.6 μg of NP (all in pcDNA3.1 vector), and 0.3 μg Pol I plasmid encoding influenza virus segment 5 vRNA or cRNA. TRIzol reagent was used for extracting total cellular RNA at 18 hours post-transfection, followed by RT and qPCR performed as described above. The primers used for RT and qPCR are detailed in Table S2.

### Split Luciferase Assay

The interaction between ANP32 proteins and the viral polymerase complex, as well as the formation of symmetric polymerase dimers, were examined using a luciferase complementation assay(36–38). In this assay, Gaussia luciferase (Gluc) is split into two inactive fragments (Gluc-N and Gluc-C) that are separately fused to target proteins A and B. Interaction between A and B brings the two Gluc fragments into proximity, allowing conformational refolding and restoration of luciferase activity. The resulting bioluminescent signal is quantifiable and positively correlates with interaction efficiency.

To determine the impact of PA mutations on the viral polymerase complex-ANP32A/B interaction, HEK293T cells in 12-well plates and co-transfected with 0.5 μg each of PB1-Gluc-N, ANP32A-Gluc-C or ANP32B-Gluc-C, PB2, as well as wild-type PA or PA-K605A/K605R/K609A mutant plasmids. At 24 h post-transfection, cells were lysed with 250 μL Renilla lysis buffer (Promega) on ice for 30 min. Gaussia luciferase activity was assayed using a Renilla luciferase detection kit (Promega) on a TECAN SPARK microplate reader, in accordance with the manufacturer’s recommended protocol.

To evaluate the impact of PA mutations on viral polymerase symmetric dimerization, HEK293T cells in 12-well plates were co-transfected with 0.5 μg each of PB2 and wild-type PA or PA-K605A/K605R/K609A mutant, along with 0.25 μg each of PB1-Gluc-N and PB1-Gluc-C. Following 24 h of additional culture, Gaussia luciferase activity was measured using the protocol described above.

### Immunoprecipitation and immunoblotting

Cells for immunoprecipitation or co-immunoprecipitation assays were seeded in 10-cm cell culture dishes prior to experimental treatment. For exogenous detection of viral protein lactylation, HEK293T cells were transfected with 10 μg empty vector or FLAG-tagged viral protein expression plasmids. For endogenous detection of viral protein lactylation, A549 cells were infected with influenza A virus at a MOI of 1 or 0.01. To detect viral PA-ATAT1/SIRT1 interaction, HEK293T cells were co-transfected with 5 μg each of FLAG-tagged PA and HA-tagged ATAT1, SIRT1 or SIRT1-ΔCata. To evaluate the impact of PA mutations on the polymerase complex-ANP32A/B interaction, HEK293T cells were transfected with 2.5 μg each of FLAG-tagged ANP32A/ANP32B, pcDNA3.1-PB1, pcDNA3.1-PB2 and pcDNA3.1-WT or PA-K605A/K605R/K609A mutants. At indicated times post-transfection or post-infection, cells were lysed with 1 mL Western and IP lysis buffer (P0013; Beyotime, China) for 30 min on ice. Cell lysates were then incubated with the indicated primary antibody-conjugated Protein A/G magnetic beads (22204; Beaver Beads, China) for 4 h at 4 ℃ with gentle rotation. After incubation, the magnetic beads were isolated using a magnetic separator and washed five times with PBST (PBS containing 0.5% Tween-20). Immunoprecipitated proteins were resolved by SDS-PAGE and then transferred onto NC membranes. The membranes were blocked with 5% non-fat milk in TBST (Tris-buffered saline containing 0.1% Tween-20) for 1 h at room temperature, followed by incubation with the indicated primary antibodies overnight at 4 ℃. After three washes with TBST, the membranes were incubated with horseradish peroxidase (HRP)-conjugated secondary antibodies for 1 h at room temperature. The protein bands were visualized using an enhanced chemiluminescence (ECL) detection system (K-12045; Advansta, USA).

### Determination of viral pathogenicity

For pathogenicity determination, female BALB/c mice (18-20 g body weight, 6-8 weeks old) were lightly anesthetized by ether inhalation and intranasally inoculated with either PBS, 10 PFU of wild-type PR8 influenza virus, or PR8 mutants carrying PA K605A, PA K605R, or PA K609A substitutions (18 mice per group). Twelve mice from each group were randomly assigned to the survival and body weight monitoring cohort and observed for 14 consecutive days, with a 30% body weight reduction considered death. At 3 and 5 d post-inoculation, three mice were randomly selected from each group, humanely euthanized, and lung tissues were harvested and split into two aliquots. One aliquot was fixed for histopathological sectioning and subsequent hematoxylin-eosin (H&E) and immunofluorescence staining; the other aliquot was homogenized, and the resulting supernatant was collected via centrifugation for viral titer quantification using TCID_50_ assays on MDCK cells.

### Statistical analyses

GraphPad Prism v9.x software was used for data collating, analyzing and graph plotting. Data are presented as the means ± standard deviation (SD) and are repeated for three times. Comparisons between two groups were performed using unpaired Student’s t-tests. Survival curves were generated using Kaplan-Meier method and analyzed via the Log-rank (Mantel-Cox) test. A *P* value < 0.05 was defined as statistically significant (*ns*, not significant; \**P* < 0.05; \*\**P* < 0.01; \*\*\**P* < 0.001; \*\*\*\**P* < 0.0001).

## Results

### Subunits of the influenza virus vRNP complex are lactylated

Influenza virus infection disrupts host cell glycolysis, leading to elevated intracellular lactate levels (26–28). We first assessed the effect of the lactate dehydrogenase A (LDHA) inhibitor oxamate on influenza virus replication. Our results showed that oxamate suppressed viral replication (Fig. 1A; Fig. S1A). Similarly, siRNA-mediated knockdown of LDHA inhibited viral replication (Fig. 1B). In contrast, exogenous lactate supplementation increased viral titers in a dose-dependent manner (Fig. 1C; Fig. S1B). Given that lactate is the key substrate for lactylation, it is hypothesized that lactate regulate viral replication through lactylation of viral proteins. Therefore, lactylation of viral vRNP subunits, the minimal replication unit of influenza virus, was investigated. Exogenous and endogenous IP assays using FLAG-antibody or pan-Lactyl Lysine antibody showed that subunits of the viral vRNP complex-PB1, PB2, PA, and NP-undergo lactylation, with PA and PB2 displaying relatively higher lactylation levels (Fig. 1D-E). In addition, exogenous sodium lactate treatment enhanced the lactylation of PB1, PB2, PA and NP proteins under both infection and transfection conditions (Fig. 1F; Fig. S1C-F). In contrast, treatment with Galloflavin, another LDHA inhibitor, decreased the lactylation levels of these viral proteins (Fig. 1G). These results indicate that lactate promotes influenza virus replication and that viral PB1, PB2, PA, and NP proteins are subjected to lactylation. To confirm lactylation of viral proteins, we concentrated viral particles, and enriched lactylated proteins using the pan-Lactyl Lysine antibody, followed by MS identification (Fig. 1H). Silver staining confirmed effective enrichment of viral proteins by the pan-Lactyl Lysine antibody, and MS analysis verified lactylation of viral PB1, PB2, PA and NP proteins, with six lactylation residues identified on the PA protein (Fig. 1I; Table S1). On this basis, subsequent studies were focused on in-depth characterization of viral PA lactylation.

**Figure 1.**
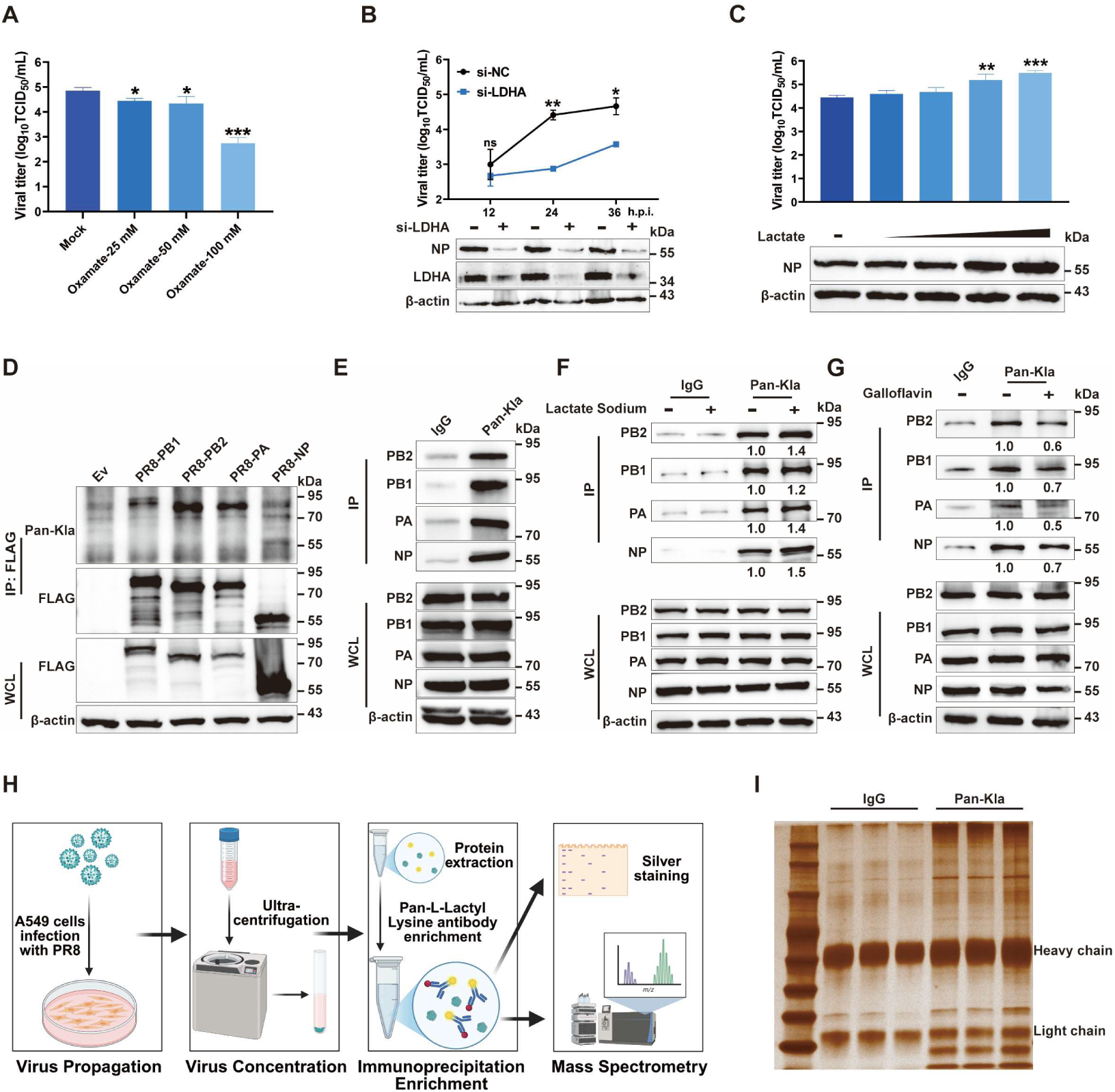
Influenza virus vRNP subunits are lactylated. (A) A549 cells were infected with PR8 influenza virus at a MOI of 0.01 and subsequently treated with 25, 50 or 100 mM of oxamate. Viral titers were quantified using TCID_50_ assay at 24 hpi. (B) A549 cells were transfected with non-targeting (NC) or LDHA-targeting siRNA; 24 hours post-transfection, cells were infected with PR8 (MOI=0.01). Viral titers were determined at various h post-infection using TCID_50_ assay; LDHA knockdown efficiency, and viral NP expression were assessed by immunoblotting. (C) A549 cells were infected with PR8 (MOI=0.01) and treated with 0, 2.5, 5, 10, or 20 mM of lactate. Viral titers were determined with TCID_50_ assay at 24 hpi; viral protein expression was detected by immunoblotting. (D) HEK293 cells were transfected with plasmids encoding FLAG-tagged viral PB1, PB2, PA, or NP. At 24 hours post-transfection, anti-FLAG immunoprecipitation was performed, and protein lactylation was detected by immunoblotting with a Pan-L-lactyl lysine antibody. (E) A549 cells were infected with PR8 (MOI=0.01); at 24 hpi, anti-Pan-L-lactyl lysine immunoprecipitation was conducted, and enrichment of viral proteins was detected by immunoblotting with viral protein-specific antibodies. (F) A549 cells were infected with PR8 (MOI=1) and treated with 20 mM sodium lactate. At 12 hpi, anti-Pan-L-lactyl lysine immunoprecipitation was performed, and viral protein enrichment was detected using immunoblotting with viral protein-specific antibodies. Band intensities were quantified by ImageJ and normalized to that of Mock-treated group. (G) A549 cells were infected with PR8 (MOI=1) and then treated with 40 μM galloflavin; anti-Pan-L-lactyl lysine immunoprecipitation was carried out at 12 hpi, and viral protein enrichment was detected by immunoblotting with viral protein-specific antibodies. Band intensities were quantified by ImageJ and normalized to that of Mock-treated group. (H-I) Workflow for the identification of influenza virus proteins by mass spectrometry following Pan-L-lactyl lysine antibody enrichment; enriched samples in (H) were subjected to silver staining (I). Schematic diagrams were generated using BioRender. Data are presented shown as mean ± SD, and are representative of three independent experiments. Statistical analyses were performed using unpaired Student’s t-tests; *ns*, not significant; \**P*<0.05, \*\**P*<0.01, \*\*\**P*<0.001.

### ATAT1 promotes influenza virus PA lactylation

During lactylation, lactyl-CoA, derived from lactate, forms covalent linkages with lysine residues on substrate proteins, a process catalyzed by enzymes. It has been shown that cellular alanine-tRNA ligases (AARS1 and AARS2) act as lactyltransferase to mediate lactylation of multiple cellular proteins (39, 40). Additionally, several acetyltransferases, including N-acetyltransferase ESCO1, ESCO2, α-tubulin N-acetyltransferase 1 (ATAT1) and histone acetyltransferase MYST1, also exert lactyltransferase activity(24, 41). Having confirmed viral PA protein is lactylated, the key enzyme mediating PA lactylation was subsequently investigated. Co-IP screening indicated that the influenza viral PA protein interacts with ATAT1 (Fig. S2A). To further verify this interaction, exogenous co-IP assays were performed using anti-FLAG or anti-HA antibodies, which confirmed the PA-ATAT1 interaction (Fig. 2A-B). Additionally, co-localization of ATAT1 and viral PA was observed under fluorescence microscopy (Fig. S2B). We next investigated the effect of ATAT1 on PA lactylation. The results showed that overexpression of ATAT1 significantly increased PA lactylation (Fig. 2C). In contrast, siRNA-mediated silencing of endogenous ATAT1 led to a notable decrease in PA lactylation level (Fig. 2D). The impact of ATAT1 on influenza virus replication was subsequently evaluated. Results showed that ATAT1 knockdown led to decreased viral titers; whereas ATAT1 overexpression promoted viral replication (Fig. 2E; Fig. S2C). The influenza virus PA protein assembles into a heterotrimeric polymerase complex with PB1 and PB2 to exert transcription and genome replication activity (42). Subsequently, the effects of ATAT1 on viral polymerase activity were assessed. ATAT1 overexpression increased viral polymerase activity in a dose-dependent manner, while ATAT1 knockdown reduced it (Fig. 2F; Fig. S2D). To further verify that ATAT1 mediates lactylation of viral PA, a ATAT1-D157N mutant, that abrogates the acetyltransferase activity of ATAT1, was constructed (43). Given the mechanistic similarity between acetylation and lactylation, we hypothesized that ATAT1-D157N also lacks lactyltransferase activity. We found that ATAT1-D157N overexpression failed to promote PA lactylation (Fig. 2G). Furthermore, compared with wild-type (WT) ATAT1, ATAT1-D157N mutant exhibited a significantly reduced ability to enhance viral polymerase activity (Fig. 2H). Collectively, our results confirm that ATAT1 facilitates lactylation of influenza virus PA protein and promotes viral replication.

**Figure 2.**
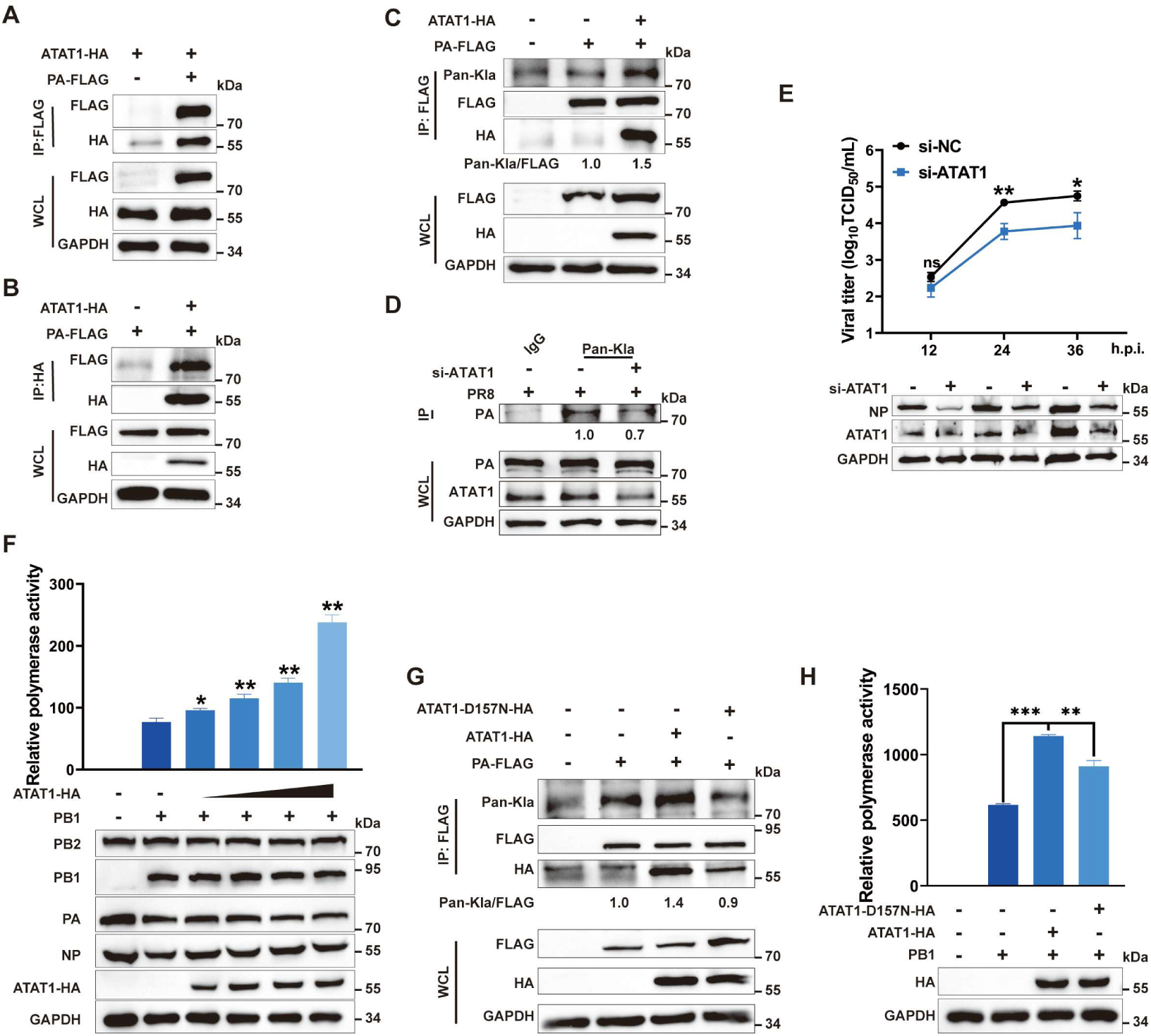
ATAT1 promotes influenza virus PA lactylation. (A-B) HEK293T cells were co-transfected with Flag-tagged PA and HA-tagged ATAT1 or empty vector; anti-FLAG (A) or anti-HA (B) co-IP assays were performed at 24 h post-transfection, co-precipitated proteins were detected by immunoblotting. (C) HEK293T cells were co-transfected with FLAG-tagged PA and either HA empty vector or HA-tagged ATAT1. At 24 h post-transfection, FLAG-tagged proteins were enriched by anti-FLAG immunoprecipitation, and PA lactylation was detected by immunoblotting with a Pan-L-lactyl lysine antibody; the band intensities were analyzed using ImageJ and normalized to empty vector-transfected group. (D) A549 cells were transfected with NC or ATAT1-targeting siRNA for 24 h, followed by influenza virus infection (MOI=1) for 12 h; lactylated proteins were enriched by a Pan-L-lactyl lysine antibody, PA enrichment was detected by immunoblotting using viral PA-specific antibodies. Band intensities were analyzed using ImageJ and normalized to NC-transfected group. (E) A549 cells were transfected with NC or ATAT1-targeting siRNA; 24 hours later, cells were infected with influenza virus at a MOI of 0.01. Viral titers were detected using TCID_50_ assay at various h post-infection; ATAT1 knockdown efficiency, and viral NP expression were assessed by immunoblotting. (F) HEK293T cells were transfected with influenza virus PA/PB1/PB2/NP, pPol I-luc reporter, pRL-TK internal control, and 0, 0.25, 0.5, 1, or 2 μg of HA-tagged ATAT1. At 24 hours post-transfection, Firefly and Renilla luciferase activities were measured using a dual-luciferase reporter assay system, and protein expression was detected by immunoblotting. (G) HEK293T cells were co-transfected with FLAG-tagged PA and HA empty vector, HA-tagged ATAT1, or HA-tagged ATAT1-D157N. At 24 h post-transfection, FLAG-tagged proteins were enriched with an anti-FLAG antibody, and PA lactylation was detected by immunoblotting using a Pan-L-lactyl lysine antibody. Band intensities were quantified by ImageJ and normalized to that of empty vector group. (H) HEK293T cells were transfected with influenza virus PA/PB1/PB2/NP, pPol I-luc reporter, pRL-TK internal control, and either HA-tagged ATAT1 or HA-tagged ATAT1-D157N. At 24 hours post-transfection, Firefly and Renilla luciferase activities were measured using a dual-luciferase reporter assay system, and protein expression was detected by immunoblotting. Data are presented as mean ± SD and are representative of three independent experiments. Statistical analyses were performed using unpaired Student’s t-tests; *ns*, not significant; \**P*<0.05, \*\**P*<0.01, \*\*\**P*<0.001.

### SIRT1 mediates de-lactylation of influenza virus PA

Like other PTMs, lactylation is a reversible process. Although no specific de-lactyltransferase has been identified to date, several de-acetyltransferases, including sirtuins 1-7 (SIRT1-7) and histone de-acetyltransferases 1-3 (HDAC1-3), have been reported to possess de-lactyltransferase activity (44–47). To screen the key enzyme mediating PA de-lactylation, we examined interactions between the afore-mentioned de-acetylases (SIRT3 and SIRT4 were excluded due to their mitochondrial localization) and viral PA. Co-IP screening indicated PA interacts strongly with SIRT1, which was further validated by both exogenous and endogenous co-IP assays (Fig. S3A; Fig. 3A-C). Next, we examined the effect of SIRT1 on PA lactylation; the results showed that overexpression of SIRT1 significantly reduced the lactylation level of PA (Fig. 3D). In contrast, siRNA-mediated silencing of SIRT1 enhanced viral PA lactylation (Fig. 3E). The impact of SIRT1 on influenza virus replication was subsequently evaluated. Notably, siRNA-mediated silencing of SIRT1 significantly promoted, whereas exogenous SIRT1 overexpression inhibited influenza virus replication (Fig. 3F; Fig. S3B). Furthermore, we demonstrated that exogenous SIRT1 overexpression inhibited viral polymerase activity in a dose-dependent manner, while knockdown of SIRT1 led to an increase in viral polymerase activity (Fig. 3G; Fig. S3C). SIRT1 contains a catalytic domain (aa 234-510) and a C-terminal regulatory segment (aa 641-665) (48). To further explore the role of these domains, a catalytic domain-deleted SIRT1 mutant (SIRT1-ΔCata) was constructed and its effect on PA lactylation was evaluated (Fig. 3H). The results showed that overexpression of SIRT1-ΔCata failed to down-regulate PA lactylation level (Fig. 3I). Meanwhile, PA failed to co-precipitate with SIRT1-ΔCata, suggesting that the catalytic domain of SIRT1 is critical for its interaction with viral PA (Fig. 3J). To further verify that SIRT1 regulates influenza virus replication via its de-lactyltransferase activity, the effect of SIRT1-ΔCata on viral proliferation was assessed. While WT SIRT1 overexpression significantly reduced viral titers, SIRT1-ΔCata failed to inhibit viral replication (Fig. 3K). Moreover, SIRT1-ΔCata lost the inhibitory effect on viral polymerase activity (Fig. 3L). Collectively, these results confirm that SIRT1 mediates de-lactylation of the viral PA, and its catalytic domain plays a critical role in regulating PA de-lactylation and restricting influenza virus replication.

**Figure 3.**
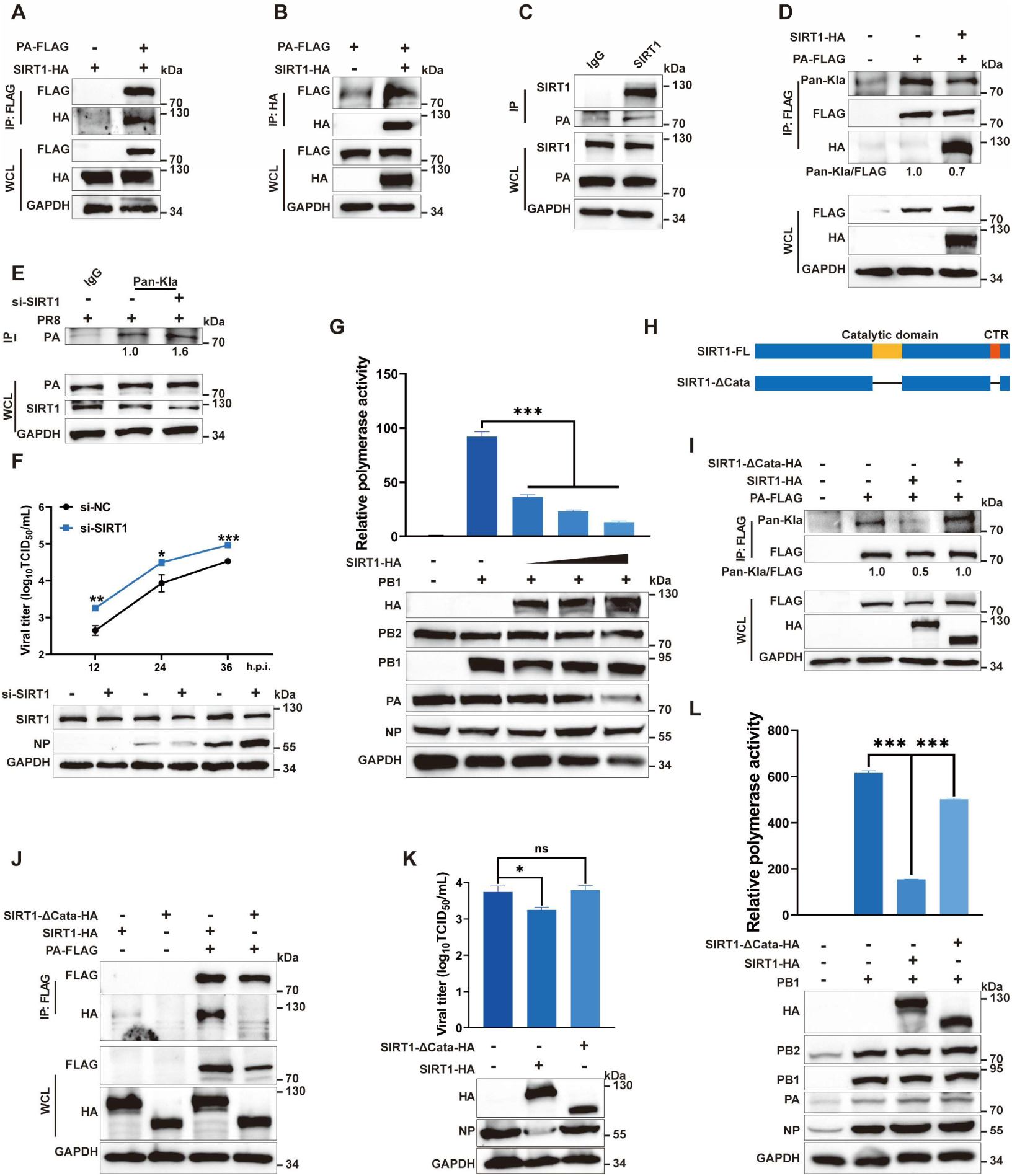
SIRT1 de-lactylates viral PA. (A-B) HEK293T cells were co-transfected with Flag-tagged PA and HA-tagged SIRT1 or empty vector; anti-FLAG (A) or anti-HA (B) co-IP assays were performed at 24 h post-transfection, co-precipitated proteins were detected by immunoblotting. (C) A549 cells were infected with influenza virus (MOI=0.01); anti-SIRT1 co-IP was performed at 24 hpi, precipitation of viral PA protein was detected by immunoblotting with viral PA-specific antibody. (D) HEK293T cells were co-transfected with FLAG-tagged PA and HA empty vector or HA-tagged SIRT1. FLAG-tagged proteins were enriched by anti-FLAG IP at 24 h post-transfection, and PA lactylation was detected by immunoblotting with a Pan-L-lactyl lysine antibody; band intensities were quantified by ImageJ and normalized to that of empty vector group. (E) A549 cells were transfected with NC or SIRT1-targeting siRNA for 24 h, followed by influenza virus infection (MOI=1) for 12 h; lactylated proteins were enriched by a Pan-L-lactyl lysine antibody, viral PA enrichment was detected by immunoblotting using viral PA-specific antibodies; band intensities were analyzed using ImageJ and normalized to NC-transfected group. (F) A549 cells were transfected with NC or SIRT1-targeting siRNA; 24 hours later, cells were infected with influenza virus at a MOI of 0.01. Viral titers were determined by TCID_50_ assay at various h post-infection; SIRT1 knockdown efficiency, and viral NP expression were detected by immunoblotting. (G) HEK293T cells were transfected with influenza virus PA/PB1/PB2/NP, pPol I-luc reporter, pRL-TK internal control, and increasing concentrations of HA-tagged SIRT1. At 24 h post-transfection, Firefly and Renilla luciferase activities were measured using a dual-luciferase reporter assay system, and protein expression was detected by immunoblotting. (H) Schematic diagram of the SIRT1 catalytic domain and C-terminal regulatory segment. (I) HEK293T cells were co-transfected with FLAG-tagged PA and HA empty vector, HA-tagged SIRT1, or HA-tagged SIRT1-ΔCata. FLAG-tagged proteins were enriched by anti-FLAG immunoprecipitation at 24 h post-transfection, and PA lactylation was detected by immunoblotting with a Pan-L-lactyl lysine antibody. Band intensities were quantified by ImageJ and normalized to that of empty vector group. (J) HEK293T cells were co-transfected with FLAG-tagged PA and HA-tagged SIRT1 or SIRT1-ΔCata. Anti-FLAG co-IP was performed at 24 h post-transfection; co-precipitated proteins were detected by immunoblotting. (K) A549 cells were transfected with HA-tagged SIRT1 or SIRT1-ΔCata; 24 hours later, cells were infected with influenza virus at a MOI of 0.01. Viral titers were determined by TCID_50_ assay at 24 hpi, protein expression was detected by immunoblotting. (L) HEK293T cells were transfected with influenza virus PA/PB1/PB2/NP, pPol I-luc reporter, pRL-TK internal control, and HA-tagged SIRT1 or SIRT1-ΔCata. Firefly and Renilla luciferase activities were measured with a dual-luciferase reporter assay system at 24 h post-transfection, and plasmid expression was detected by immunoblotting. Data are presented as mean ± SD and are representative of three independent experiments. Statistical analyses were performed using unpaired Student’s t-tests; *ns*, not significant; \**P*<0.05, \*\**P*<0.01, \*\*\**P*<0.001.

### K605/K609 residues on PA are critical for influenza virus replication and pathogenicity

As MS analysis identified six potential lactylation residues on viral PA protein, we next evaluated the biological impacts of these residues on influenza virus replication and pathogenicity. Conservation analysis revealed that all residues except K356 on viral PA are highly conserved among influenza viruses from different host species (Fig. S4A). Spatial location analysis revealed surface accessibility of these residues within the vRNA-bound polymerase complex (Fig. S4B). Subsequent site-directed mutagenesis coupled with IP analysis revealed that substitution mutations at K356, K358, K361 and K362 moderately attenuated PA lactylation, while mutations at K605 and K609 strong reduced PA lactylation (Fig. 4A). Unlike macromolecular modifications such as ADP-ribosylation, lactylation modulates protein function most likely by altering the charge state of lysine residues to modify its hydrogen bonding capacity (49). Therefore, the afore-mentioned residues were mutated to either alanine (electrically neutral) or arginine (positively charged), and their effects on viral polymerase activity were examined. Results showed that PA K605A/K605R and K609A mutations significantly reduced viral polymerase activity, suggesting that the charge state of PA K605 requires dynamic regulation, while K609 need to maintain a constant positive charge state (Fig. 4B). To determine the specific effect of PA K605/K609 mutations on the transcriptional or replicative function of the polymerase, we employed a polymerase activity complementation system: the PA C95A mutant retains only transcriptional activity, the PA D108A mutant retains only replicative activity, and co-expression of these two mutants restores polymerase activity (Fig. S4C) (50). Complementation assays showed that co-expression of PA K605/K609 mutants with PA C95A did not restore polymerase activity, whereas co-expression with PA D108A restored polymerase activity to that of WT PA (Fig. 4C). These results suggest PA K605/K609 are critical for the genomic replication function of viral polymerase complex.

**Figure 4.**
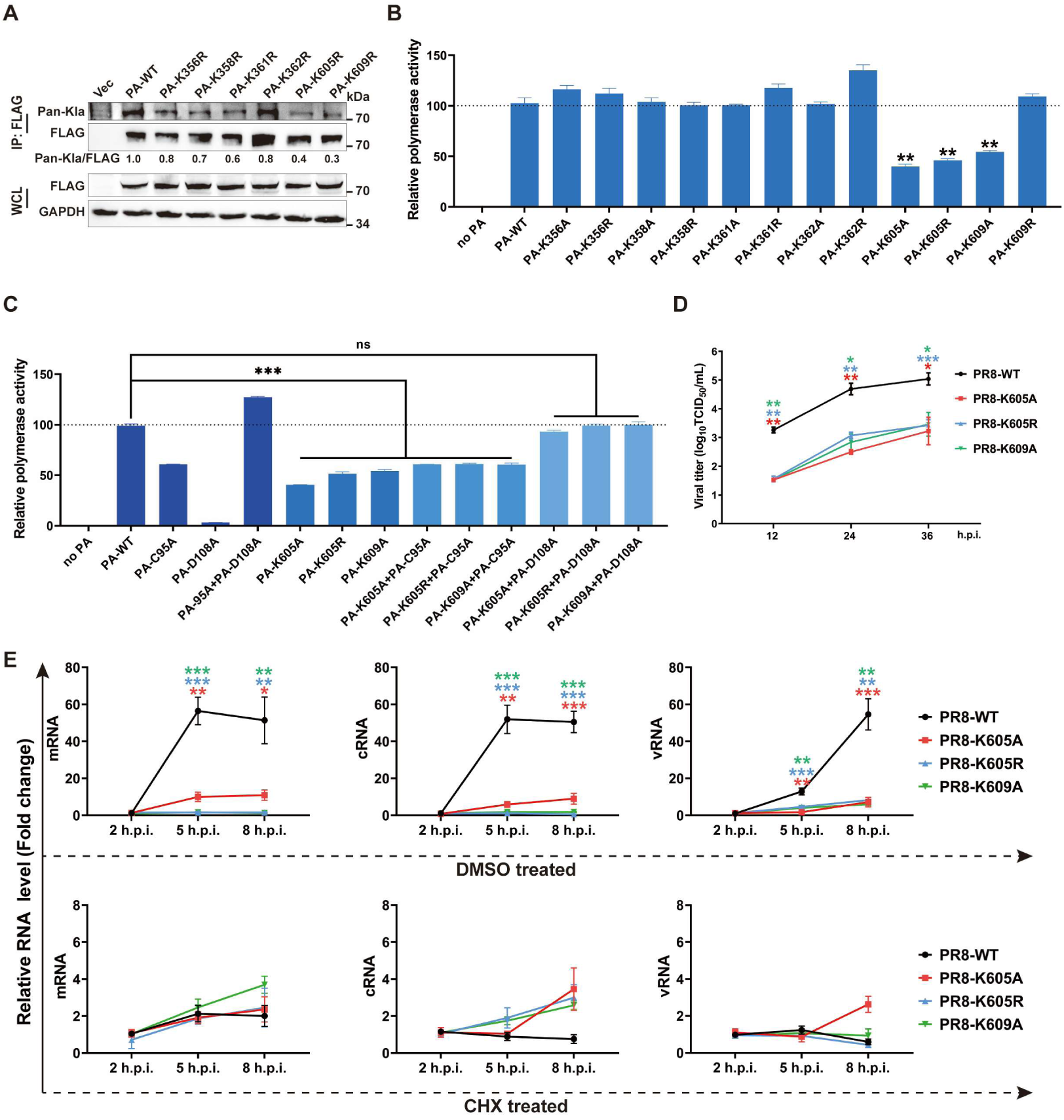
K605/K609 residues of PA are critical for influenza virus replication. (A) HEK293T cells were transfected with FLAG-tagged PA-WT or PA-K356R/K358R/K361R/K362R/K605R/K609R mutants; anti-FLAG IP was performed at 24 h post-transfection, and PA lactylation was detected by immunoblotting with a Pan-L-lactyl lysine antibody. Band intensities were quantified by ImageJ and normalized to that of WT-PA-transfected group. (B) HEK293T cells were transfected with influenza virus PB1/PB2/ NP, pPol I-luc reporter, pRL-TK internal control, and either WT PA or PA carrying alanine- or arginine-substituted mutations at residues 356, 358, 361, 362, 605, or 609. At 24 hours post-transfection, Firefly and Renilla luciferase activities were measured with a dual-luciferase reporter assay system. (C) HEK293T cells were transfected with influenza virus PB1/PB2/NP, pPol I-luc reporter, pRL-TK internal control, PA-K605A/K605R/K609A mutants, and PA-C95A or PA-D108A mutant; Firefly and Renilla luciferase activities were measured at 24 h post-transfection with a dual-luciferase reporter assay system. (D) A549 cells were infected with WT or PA K605A/K605R/K609A mutant PR8 (MOI=0.01); viral titers in the supernatant were determined at 12/24/36 h post-infection by TCID_50_ assay. (E) A549 cells were infected with WT or PA K605A/K605R/K609A mutant PR8 (MOI=1), treated with DMSO or 100 μg/mL CHX; total cellular RNA was extracted at 2/5/8 h post-infection, and the fold-change of viral m/c/vRNA was detected by qRT-PCR with GAPDH serving as the internal control. Data are presented as mean ± SD and are representative of three independent experiments. Statistical analyses were performed using unpaired Student’s t-tests; *ns*, not significant; \**P*<0.05, \*\**P*<0.01, \*\*\**P*<0.001.

To further validate the effect of PA K605/K609 mutations on viral replication, WT and mutant viruses carrying PA K605/K609 mutations were rescued using reverse genetics. Viral growth curve determination demonstrated that PA K605/K609 mutations significantly inhibited influenza virus replication (Fig. 4D). To verify the impact of PA K605/K609 substitutions on viral genome replication during infection, cells were infected with equal doses of WT or mutant viruses, and the levels of viral mRNA, cRNA and vRNA were measured at various time points post-infection. The results showed that all three species of viral RNA exhibited markedly lower levels in the mutant virus-infected group compared with the WT control (Fig. 4E). In contrast, no notable differences in viral mRNA abundance were detected between mutant and WT virus-infected groups following cycloheximide (CHX) treatment, further confirming that PA K605/K609 mutations specifically inhibit the viral genome replication process (Fig. 4E). To evaluate the impact of PA K605/K609 residues on viral pathogenicity, mice were inoculated with a lethal dose of the WT or an equivalent dose of the PA K605/K609 mutant virus, with body weight changes and survival rates monitored over 14 consecutive days. Pathological changes, alongside viral titers, in mouse lung tissues were assessed at 3 and 5 d post-infection. Results showed that mice infected with the PA K605/K609 mutant virus exhibited milder body weight loss and lung pathological changes, lower viral titers in lung tissues, smaller distribution range of viral antigens in the lungs, and a significant higher survival rate compared with mice infected with WT virus (Fig. 5A-F). Taken together, these results demonstrate that PA residues K605/K609 are critical for the replication and pathogenicity of influenza virus.

**Figure 5.**
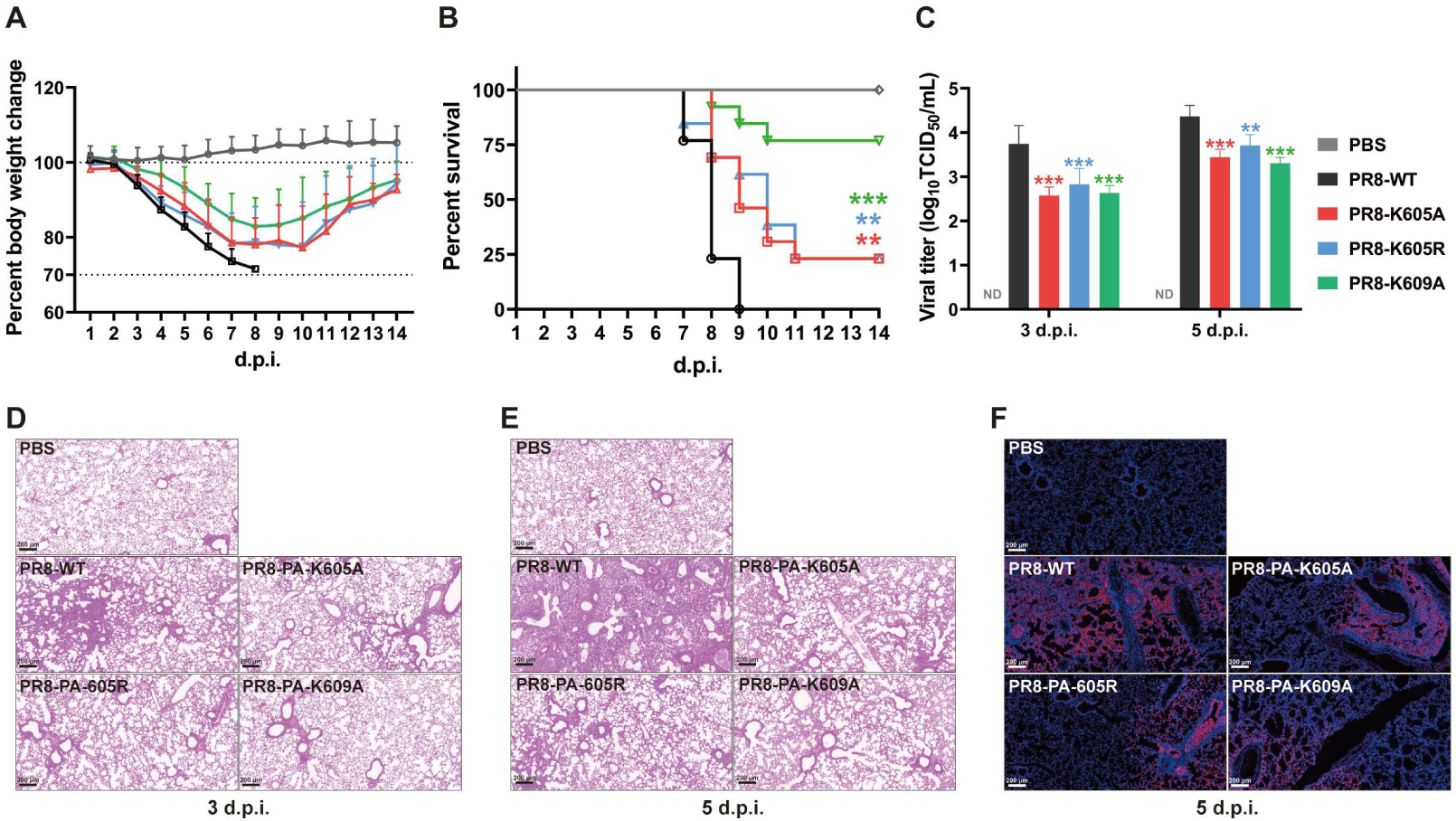
K605/K609 residues of PA are vital for influenza virus pathogenicity. Six- to eight-week-old female BALB/c mice were intranasally infected with a lethal dose (10 PFU) of WT or PA-K605A/K605R/K609A mutant PR8. Body weight (A) and survival rate (B) were monitored for 14 consecutive days. (C) Viral titers in lung tissues were determined at 3 and 5 dpi by TCID_50_ assays. (D-E) Lung tissues were collected at 3 and 5 dpi for pathological section and hematoxylin-eosin (H&E) staining (Scale bar, 200 μm). (F) Virus distribution in the lung tissues at 5 dpi was analyzed by immunofluorescence staining (Scale bar, 200 μm). Statistical analyses were performed using unpaired Student’s t-tests for viral titers and Log-rank (Mantel-Cox) test for survival curves; \*\**P*<0.01, \*\*\**P*<0.001.

### Lactylation of PA at K605/K609 facilitates ANP32-mediated polymerase asymmetric dimerization

To initiate genomic replication, newly synthesized polymerase subunits need to translocate into the nucleus to form functional dimers. Notably, viral PA interacts with PB1 to form a complex for nuclear import (51). Co-IP assay showed that PA K605/K609 mutations did not affect the PA-PB1 interaction (Fig. S5A). During viral genome replication, the polymerase dimer exists in two major conformational states: asymmetric and symmetric dimers. The asymmetric dimer, bridged by ANP32 proteins, is critical for both cRNA and vRNA production, while the symmetric dimer generates progeny vRNAs from cRNA templates. Spatial location analysis revealed that PA K605/K609 locates at the interaction interface of the polymerase asymmetric dimer (Fig. 6A; Fig. S5B). Since ANP32 proteins are key bridging factors for the asymmetric polymerase dimer, a luciferase complementation system was employed to reflect the formation of viral polymerase asymmetric dimer by detecting the interaction between viral polymerase complex and ANP32 proteins (36). Results showed that PA K605/K609 mutations significantly inhibited the interaction between viral polymerase complex and ANP32 proteins (Fig. 6B; Fig. S5C). Meanwhile, PA K605/K609 mutations led to a marked reduction in the amount of polymerase subunits precipitated by ANP32 proteins (Fig. 6C; Fig. S5D). To exclude the interference of endogenous ANP32 proteins, the luciferase complementation assays were performed in ANP32A/B double-knockout (DKO) cells, and consistent results were obtained (Fig. S5E-F).

**Figure 6.**
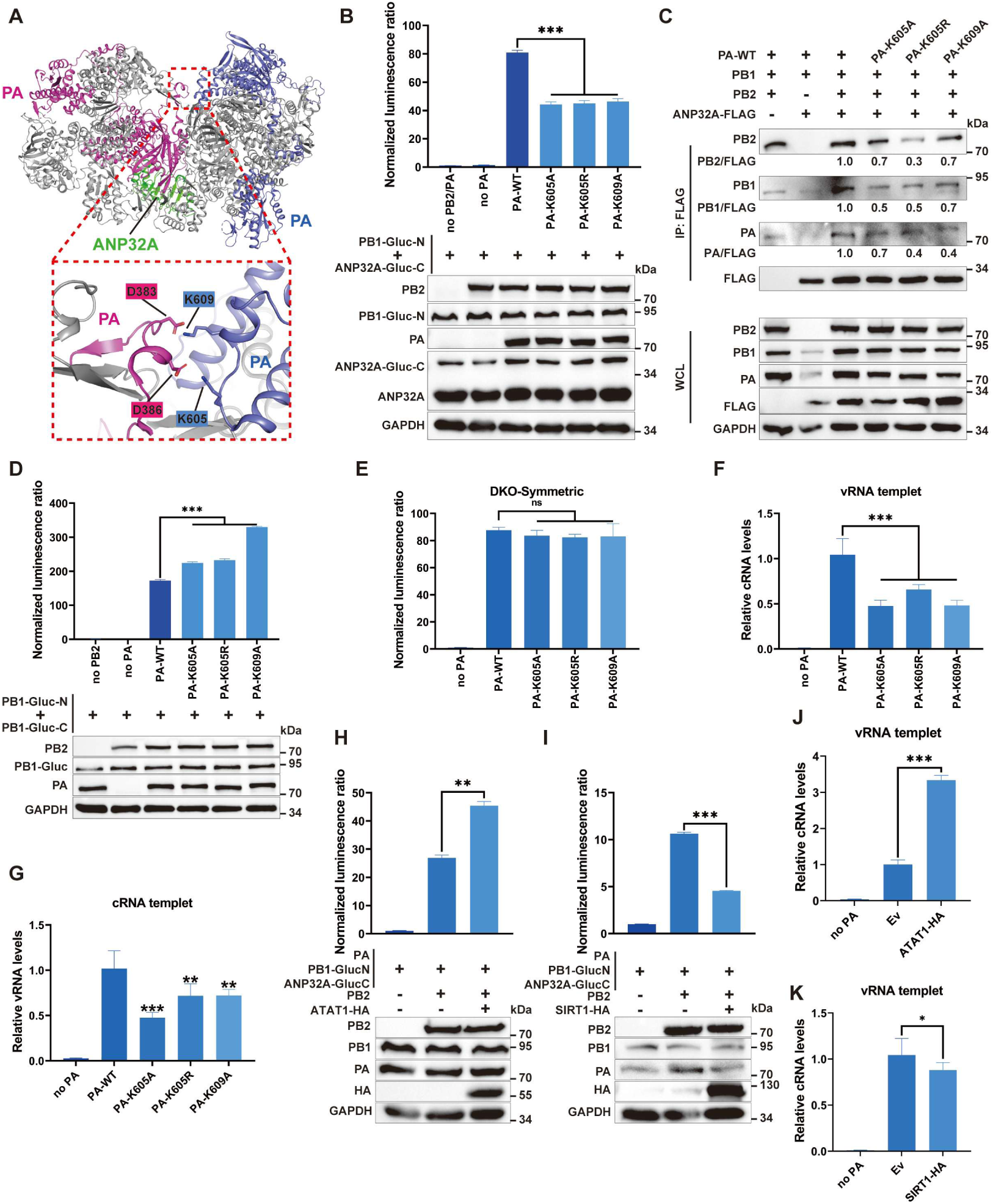
Lactylation of PA at K605/K609 facilitates ANP32A-mediated polymerase asymmetric dimerization and viral genome replication. (A) H7N9 influenza virus polymerase asymmetric dimer (PDB: 8RMR) in complex with ANP32A (green) showing PA K605 and K609. (B) HEK293T cells were co-transfected with PB2, PB1-Gluc-N, ANP32A-Gluc-C, and WT or PA K605A/K605R/K609A mutants; Gaussia luciferase activity was measured at 24 h post-transfection; protein expression was detected by immunoblotting. (C) HEK293T cells were co-transfected with FLAG-ANP32A, PB1, PB2, and WT or PA K605A/K605R/K609A mutants; anti-FLAG co-IP was performed at 24 h post-transfection; co-precipitated proteins were detected by immunoblotting with targeted antibodies. Band intensities were quantified by ImageJ and normalized to that of WT-PA-transfected group. (D) HEK293T cells were co-transfected with PB2, PB1-Gluc-N, PB1-Gluc-C, and WT or PA K605A/K605R/K609A mutants; Gaussia luciferase activity was measured at 24 h post-transfection; protein expression was detected by immunoblotting. (E) ANP32A/B double-knockout HEK293T cells were co-transfected with PB2, PB1-Gluc-N, PB1-Gluc-C, WT or PA K605A/K605R/K609A mutants; Gaussia luciferase activity was measured at 24 h post-transfection. (F-G) HEK293T cells were transfected with PB1/PB2/NP, Pol I-viral segment 5 vRNA (F) or Pol I- viral segment 5 cRNA (G), WT or PA K605A/K605R/K609A mutants; total RNA was extracted at 18 h post-transfection, and viral cRNA/vRNA levels were quantified by qPCR (GAPDH as internal control). (H-I) HEK293T cells were co-transfected with PB2, PB1-Gluc-N, ANP32A-Gluc-C, PA, and HA-ATAT1 (H) or HA-SIRT1 (I); Gaussia luciferase activity was measured at 24 h post-transfection; protein expression was detected by immunoblotting. (J-K) HEK293T cells were transfected with PA/PB1/PB2/NP, Pol I- viral segment 5 vRNA, and HA-ATAT1 (J) or HA-SIRT1 (K); total RNA was extracted at 18 h post-transfection, and viral cRNA levels were quantified by qPCR (GAPDH as internal control). Data are presented as mean ± SD and are representative of three independent experiments. Statistical analyses were performed using unpaired Student’s t-tests; *ns*, not significant; \**P*<0.05, \*\**P*<0.01, \*\*\**P*<0.001.

Further spatial analysis indicated that these residues are not positioned at the interaction interface of the viral polymerase symmetric dimer (Fig. S5G). A previous finding suggested the viral polymerase dimer maintains a conformational equilibrium (36); we therefore hypothesized that PA K605/K609 mutations may also affect the formation of the symmetric dimer. Luciferase complementation assays showed that PA K605/K609 mutations enhanced polymerase symmetric dimer formation, an effect abrogated in ANP32A/B double-knockout cells (Fig. 6D-E). To further assess the biological impacts of PA K605/K609 residues on influenza virus genomic replication, vRNA or cRNA templates were added to the vRNP reconstitution system, and downstream cRNA or vRNA levels were subsequently measured. Results showed that PA K605/K609 mutations markedly reduced cRNA and vRNA levels (Fig. 6F-G). To further confirm that the reduction in polymerase asymmetric dimerization and RNA synthesis caused by PA K605/K609 mutagenesis is attributed by the loss of lactylation at these residues, the effects of ATAT1 and SIRT1 on polymerase asymmetric dimerization were analyzed. Overexpression of ATAT1 significantly enhanced the interaction between ANP32 proteins and the viral polymerase complex, whereas SIRT1 reduced this interaction (Fig. 6H-I; Fig. S5H-I). Moreover, ATAT1 overexpression markedly increased the first step of viral genome replication-cRNA synthesis (Fig. 6J). SIRT1 overexpression partially suppressed cRNA production using vRNA as the template (Fig. 6K). Collectively, these results indicate that lactylation of viral PA protein at residues K605/K609 promotes ANP32-mediated viral polymerase asymmetric dimerization and genome replication.

## Discussion

In this study, subunits of the influenza virus vRNP complex were found to be lactylated. Recent study has also reported lactylation on subunits of the influenza virus vRNP, which is consistent with our findings (52). In addition to vRNP subunits, our MS results identified lactylation of the viral glycoprotein HA (Table S1), suggesting that lysine lactylation is a widespread PTM among influenza viral proteins and contributes critically to efficient viral replication. While MS confirmed lactylation of vRNP subunits, six lactylation residues were identified on viral PA protein, with far fewer residues detected on other subunits. This observation is presumably attributable to the low lactylation level of viral proteins and the limited sensitivity of conventional MS, which hinder the comprehensive identification of lactylated residues on viral proteins (53). In this study, mature viral particles were concentrated and lysed for mass spectrometric analysis (Fig. 1H). Although lactylation on viral proteins within mature virions is likely functionally relevant, PTMs are highly dynamic, and modification profiles of viral proteins in virions may differ substantially from those of nascent or replicating viral proteins (54). For a more complete mapping of lactylated residues across the influenza proteome, future studies may employ exogenous expression of viral proteins, induction of lactylation via lactate supplementation, and targeted mass spectrometric analysis.

The influenza virus PA protein mediates “cap-snatching” during viral transcription and plays a critical role in the assembly of the viral RNA polymerase complex (55). Several PTMs have been identified on PA, among which acetylation positively regulates its endonuclease activity and stability (56, 57). While phosphorylation of PA at serine 225 enhances the fitness of H5N1 influenza virus in mice, phosphorylation at tyrosine 393 impairs viral RNA 5’ promoter binding to the polymerase complex (58, 59). However, the full landscape of PA PTM spectrum remains largely uncharacterized. In this study, we identified multiple lactylated lysine residues on viral PA, with lactylation at K605 and K609 exhibiting functional importance. Lysine residues are susceptible to multiple PTMs, including acetylation, ubiquitination and SUMOylation, with extensive crosstalk existing among these distinct modifications (60, 61). Previous study has indicated that lysine residues K605 and K609 of PA may be subjected to ubiquitination, representing a PTM hotspot within viral PA protein (62). Our IP assay revealed that mutations at PA residues 605 and 609 did not significantly alter PA ubiquitination level, a finding likely due to the high basal level of PA ubiquitination, where single-site substitutions are insufficient to alter the global ubiquitination status (Fig. S6). Nevertheless, the functional crosstalk among PTMs at these residues in regulating PA activity warrants further investigation.

Here, we discovered that ATAT1 mediates lactylation of influenza virus PA. ATAT1 is the primary enzyme responsible for acetylating the lysine 40 residue of α-tubulin in microtubules(63). ATAT1 dependent mechanisms have also been implicated in viral replication, as ATAT1 mediated NAT10 lactylation promotes KSHV replication (24). Moreover, the Epstein-Barr virus (EBV) BHRF1 protein interacts with ATAT1 to induce microtubule hyperacetylation, which supports centrosome-associated mito-aggresome formation and proviral mitophagy, thereby enabling EBV to evade IFN induction (64). Furthermore, we found that SIRT1 mediates the de-lactylation of the viral PA protein. Consistent with this observation, recent study has confirmed that SIRT1 mediates the de-lactylation of influenza virus PA, PB2, and NP proteins, thereby inhibiting viral replication (52). SIRT1 is a NAD⁺-dependent deacetylase that modulates the acetylation status of numerous cellular and viral proteins. SIRT1 promotes antiviral innate immunity by deacetylating IRF3/7 to drive their liquid-liquid phase separation and type I interferon production (65). In addition, SIRT1 negatively regulates human T-cell leukemia virus type 1 (HTLV-1) Tax-mediated viral LTR transcription through direct interaction with Tax (66). These findings support that ATAT1 and SIRT1 are involved in the replication of diverse viruses, representing promising host directed targets for broad spectrum antiviral intervention. Consistent with this idea, pterostilbene was shown to alleviate H1N1 induced lung injury by activating the AMPKα/SIRT1/PGC1α pathway to inhibit NF-κB and p38 MAPK signaling while counteracting STAT1 activation (67). Furthermore, the SIRT1 activator SRT2183 exerts protective effects for mild IAV infection (68). Beyond influenza, it will be valuable to explore the potential of ATAT1 inhibitors and SIRT1 activators in the control of other viruses.

During genome replication, the influenza virus polymerase complex forms dimers and adopts distinct conformational states. High-resolution structural studies have defined the architecture of both asymmetric and symmetric polymerase dimers, providing key insights into the mechanisms of viral genome replication (6, 7, 69, 70). However, the molecular mechanisms by which host factors regulate the assembly and dynamics of polymerase dimers remain poorly understood. Here, we show that ATAT1-SIRT1 regulated lactylation of PA at K605/K609 promotes the assembly of the ANP32 mediated asymmetric polymerase dimer. A previous study proposed that viral polymerase dimers maintain a conformational equilibrium, yet the mechanisms that fine tune this balance remain unclear (36). Although K605/K609 are not located at the interaction interface of the symmetric polymerase dimer, we observed enhanced symmetric dimerization following PA K605/K609 mutagenesis, and this effect was abolished in ANP32A/B double knockout cells. These results suggest that PA lactylation regulate the conformational balance between polymerase asymmetric and symmetric dimers. Besides lactylation, ubiquitination of PB1 at K578 has been shown to regulate polymerase dimer switching and coordinate sequential cRNA/vRNA synthesis by disrupting PB1-PB2-N1 binding and inhibiting symmetric dimer formation (62). We thus propose that influenza virus hijack multiple host PTMs to coordinately control the dynamic equilibrium of polymerase dimers, ensuring ordered genome replication and efficient viral proliferation. However, the precise regulatory network underlying polymerase conformational dynamics during the full replication cycle remains to be further explored.

In summary, we demonstrate that lactylation of influenza virus PA, controlled by the host enzymes ATAT1 and SIRT1, promotes viral RNA synthesis during genome replication by facilitating ANP32 mediated assembly of the viral polymerase asymmetric dimer (Figure 7). These findings expand our understanding of the metabolic and post translational regulatory network underlying influenza virus infection and provide a mechanistic basis for the development of host directed antiviral strategies against influenza.

**Figure 7.**
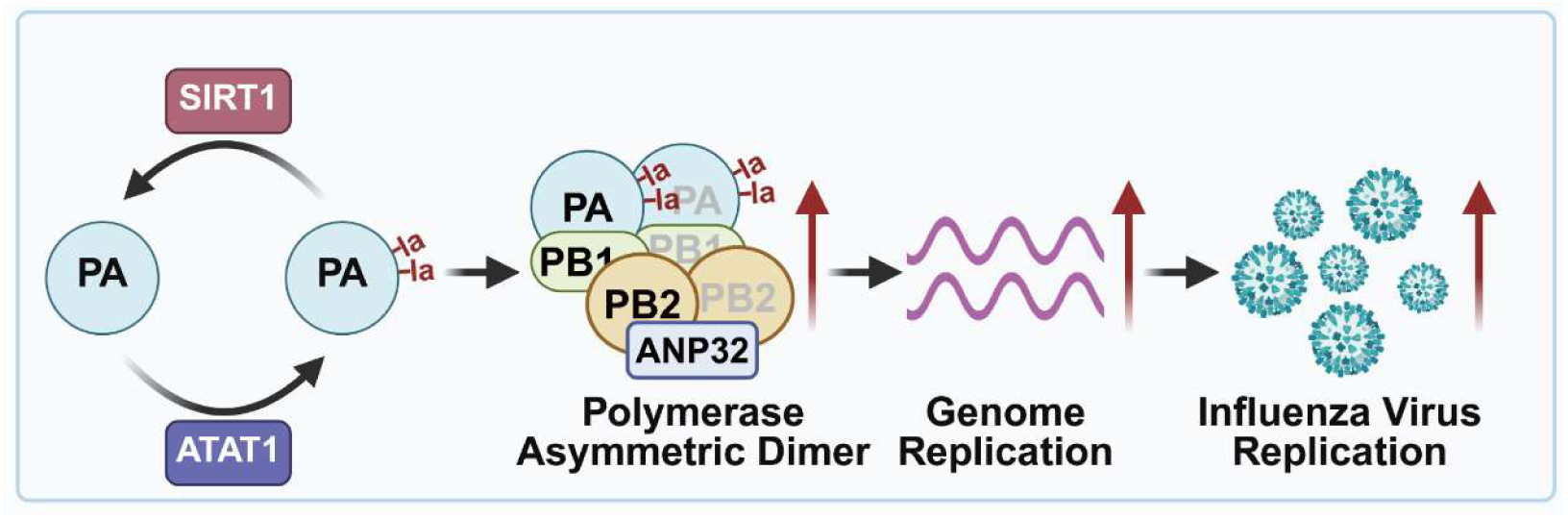
Schematic model for PA lactylation-driven viral polymerase asymmetric dimerization and influenza virus replication. ATAT1 promotes, whereas SIRT1 inhibits viral PA lactylation. Lactylation of PA at lysine residues K605/K609 facilitates the ANP32-mediated assembly of viral polymerase asymmetric dimers, which promotes viral genome replication and progeny viral production. Schematic diagram was generated using BioRender.

## Ethics declarations

Animal experiments were conducted in strict accordance with the Guidelines for the Ethical Review of Laboratory Animal Welfare and approved by the Animal Management and Ethics Committee of Huazhong Agricultural University under the approval identifier HZAUMO-2026-0057.

## Conflict of Interest

The authors declare no conflict of interest.

## Acknowledgements

This work was supported by the National Natural Science Foundation of China (32430104 to H.Z.), the National Key Research and Development Program (2024YFE0106100 to J.Z.), the Fundamental Research Fund for the Central Universities (2662025DKPY009 to H.Z.), Hubei Hongshan Laboratory (2022hszd005 to H.Z.), the earmarked fund for CARS-41 to H.Z, the Postdoctoral Fellowship Program (Grade C) of China Postdoctoral Science Foundation (GZC20251980 to T.C).

The ANP32A/B double-knockout HEK293T cell line was a kind gift from Prof. Xiaojun Wang (Harbin Veterinary Research Institute, CAAS). We are grateful to Prof. Wen Su (Huazhong Agricultural University), Pan Tao (Huazhong Agricultural University), Rui Luo (Huazhong Agricultural University), Haihong Hao (Huazhong Agricultural University) and Dr. Jun Ma (Huazhong Agricultural University) for critical discussions on experiments and manuscript drafting. We would also like to express our sincere gratitude to all staff members of the Large Instrument Open Sharing Platform of Huazhong Agricultural University for their generous support and assistance.

## Data availability

Source data are provided as supplementary source data files.

---

**Figure S1.**
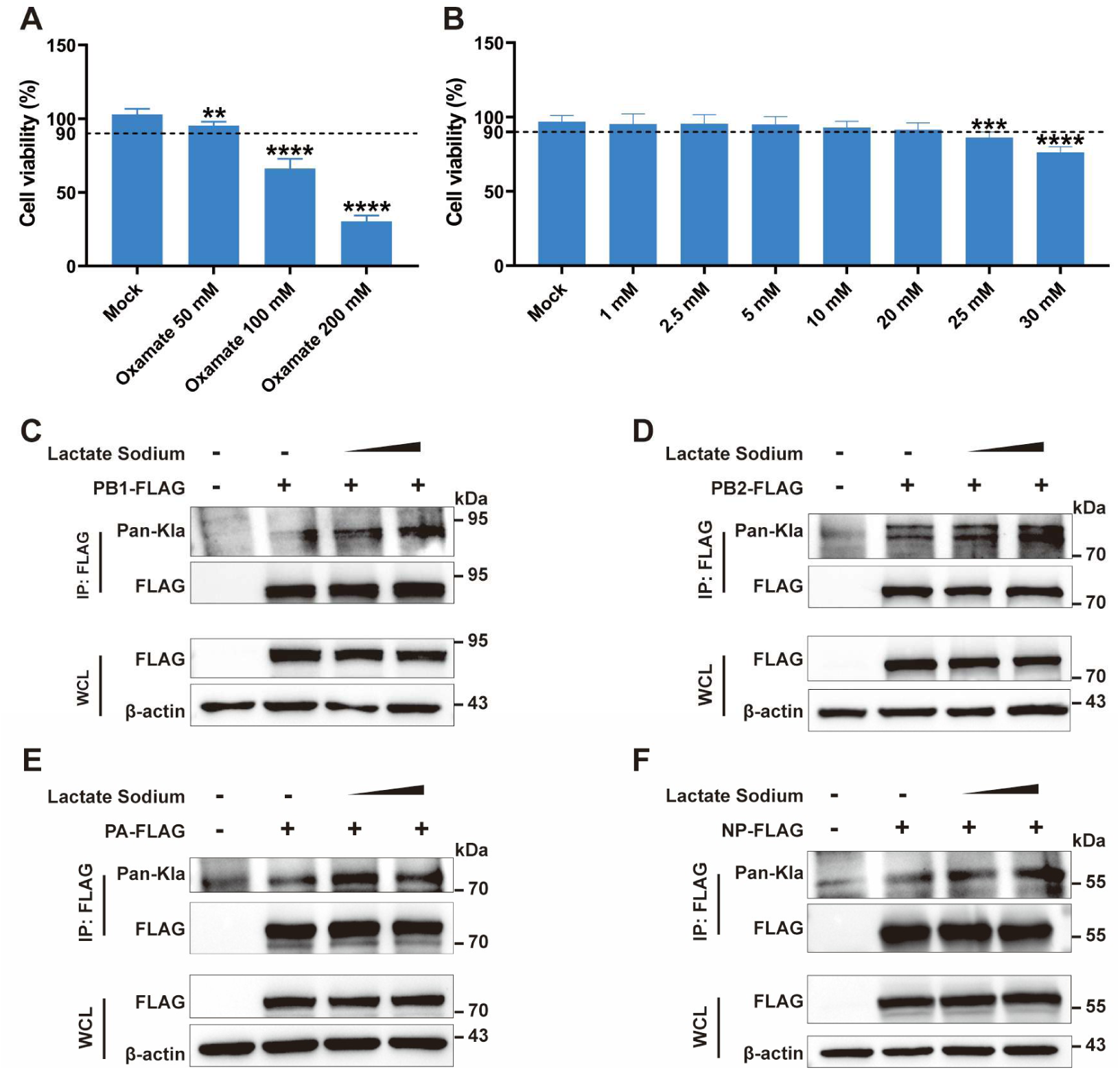
Influenza virus vRNP subunits are lactylated. (A) A549 cells were treated with 0, 50, 100, or 200 mM of oxamate for 24 h; cell viability was measured by CCK-8 assay and normalized to that of non-treated cells. (B) A549 cells were treated with 0, 1, 2,5, 5, 10, 20, 25, or 30 mM of lactate for 24 h; cell viability was measured by CCK-8 assay and normalized to that of non-treated cells. (C-F) HEK293T cells were transfected with Flag-tagged PB2 (C), PB1 (D), PA (E), or NP (F), and treated with 0, 10, or 20 mM of sodium lactate at 6 h post-transfection. Anti-Flag IP was performed at 24 h post-transfection, and immunoprecipitated samples were detected by immunoblotting with a Pan-L-lactyl lysine antibody. Data are shown as mean ± SD and are representatives of three independent assays. Statistical analyses were performed using unpaired Student’s t-tests; \*\**P* < 0.01; \*\*\**P* < 0.001; \*\*\*\**P* < 0.0001.

**Figure S2.**
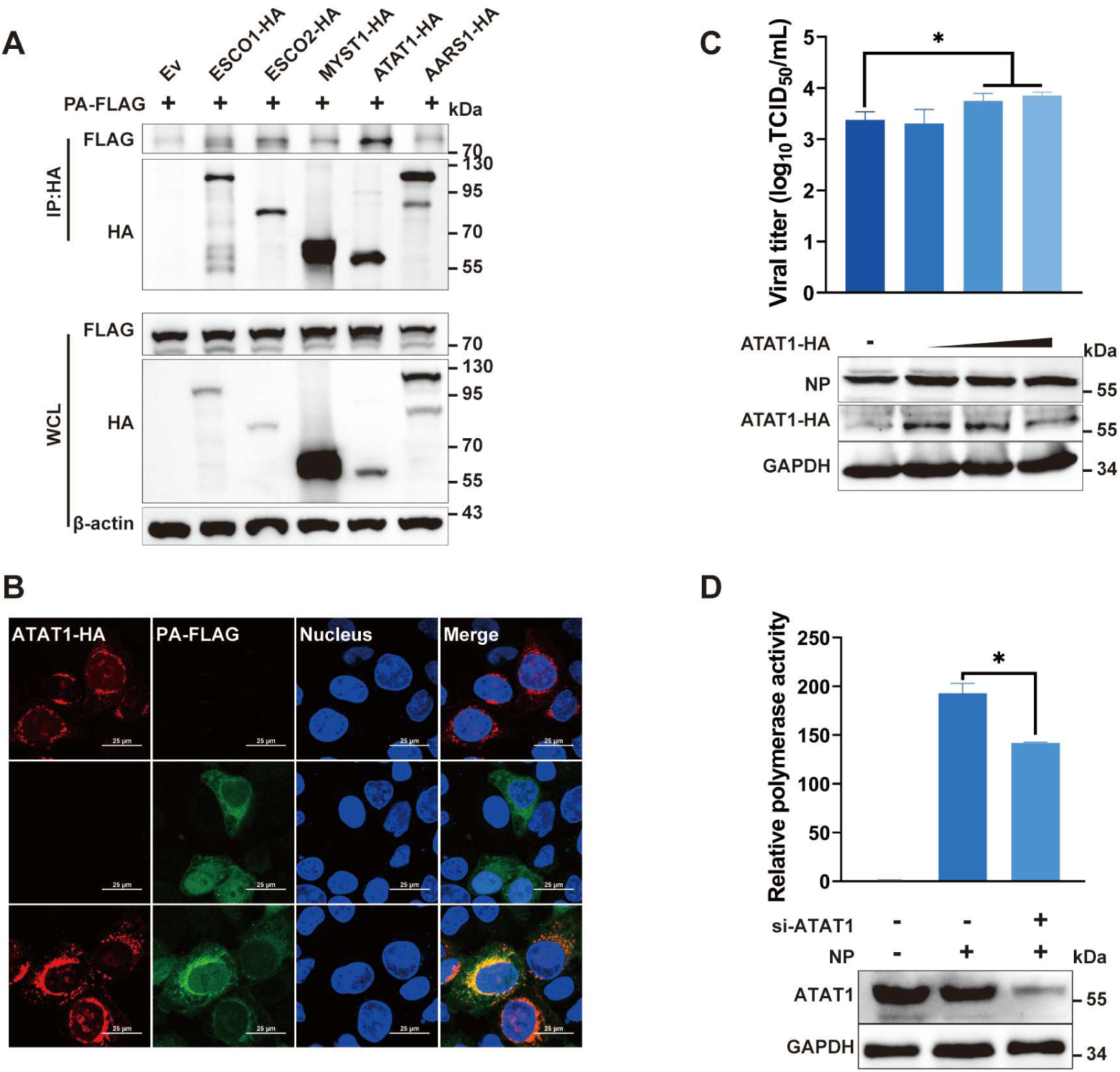
ATAT1 interacts with viral PA and promotes influenza virus replication. (A) HEK293T cells were co-transfected with FLAG-tagged PA and HA-tagged ESCO1, ESCO2, ATAT1, MYST1, AARS1, or HA empty vector. Anti-HA co-IP was performed at 24 h post-transfection; co-precipitated proteins were detected by immunoblotting. (B) HEK293T cells were single- or co-transfected with Flag-tagged PA and HA-tagged ATAT1; indirect immunofluorescence assay (IFA) was performed at 24 h post-transfection using anti-HA and anti-FLAG antibodies (Scale bar, 25 μm). (C) A549 cells were transfected with increasing doses of HA-tagged ATAT1 for 24 h, then infected with PR8 (MOI=0.01); viral titers were determined by TCID_50_ assay at 24 hpi; and protein expression levels were detected by immunoblotting. (D) HEK293T cells were transfected with NC or ATAT1-targeting siRNA, and 6 h later with influenza virus PA/PB1/PB2/NP, pPol I-luc reporter and pRL-TK internal control. Firefly and Renilla luciferase activities were measured at 24 h post-transfection using a dual-luciferase reporter assay system. Data are presented as mean ± SD and are representative of three independent experiments. Statistical analyses were performed using unpaired Student’s t-tests; \**P* < 0.05.

**Figure S3.**
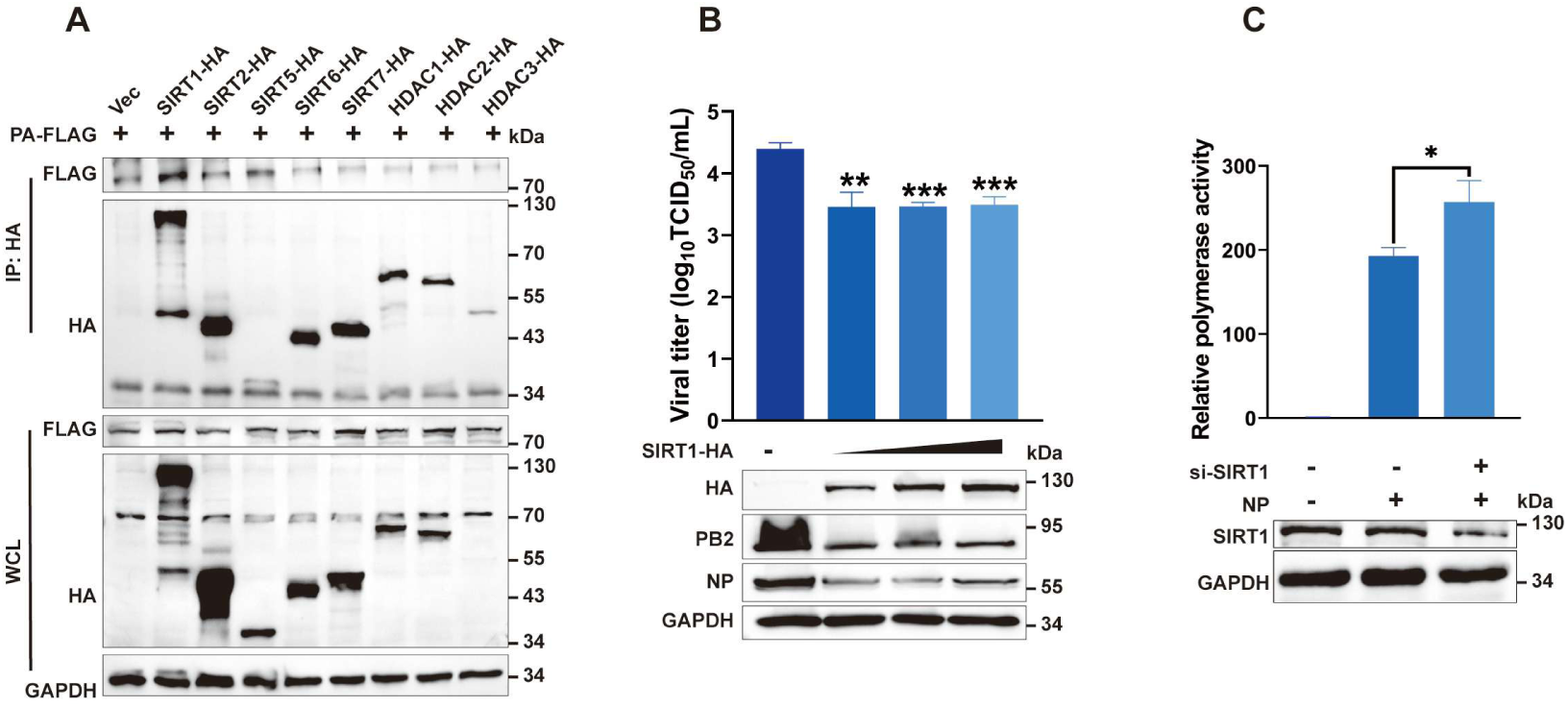
SIRT1 interacts with viral PA and inhibits influenza virus replication. (A) HEK293T cells were co-transfected with FLAG-tagged PA and HA-tagged SIRT1, SIRT2, SIRT5, SIRT6, SIRT7, HDAC1, HDAC2, HDAC3, or HA empty vector. Anti-HA co-IP was performed at 24 h post-transfection; co-precipitated proteins were detected by immunoblotting. (B) A549 cells were transfected with empty vector or increasing doses of HA-tagged SIRT1 for 24 h, then infected with PR8 (MOI=0.01); viral titers were determined by TCID_50_ assay at 24 hpi; protein expression levels were detected by immunoblotting. (C) HEK293T cells were transfected with NC or SIRT1-targeting siRNA, and 6 h later with influenza virus PA/PB1/PB2/NP, pPol I-luc reporter and pRL-TK internal control; Firefly and Renilla luciferase activities were measured at 24 h post-transfection using a dual luciferase reporter assay system. Data are presented as mean ± SD and are representative of three independent experiments. Statistical analyses were performed using unpaired Student’s t-tests; \**P* < 0.05, \*\**P* < 0.01, \*\*\**P* < 0.001.

**Figure S4.**
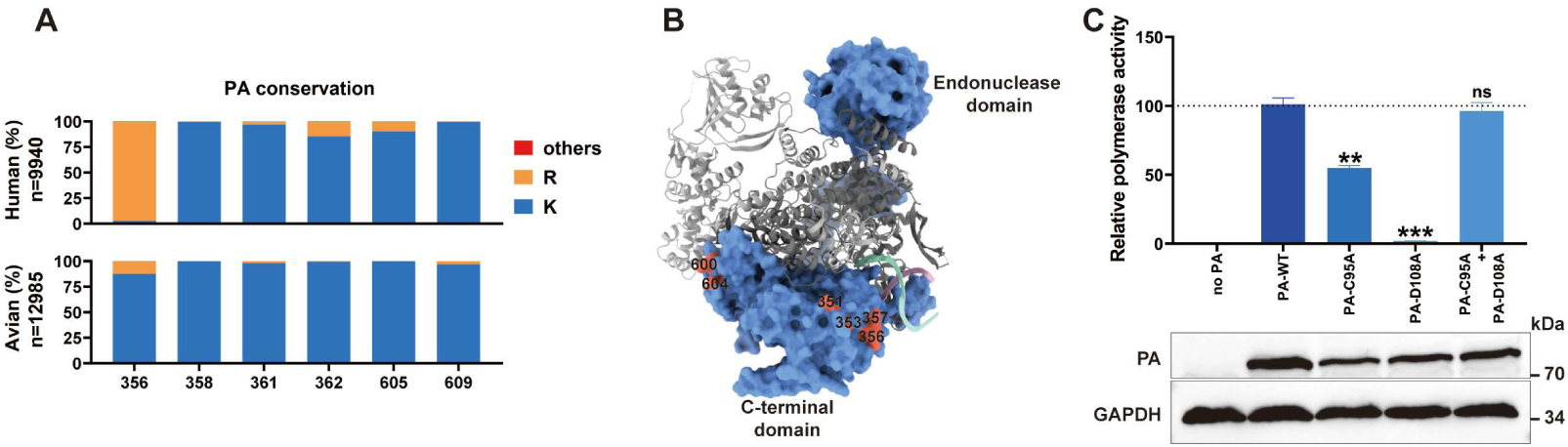
K605/K609 residues of PA are critical for influenza virus replication. (A) PA sequences from human (n = 9940), and avian (n = 12985) influenza A virus isolates were downloaded from the NCBI Influenza Database (as of September 10, 2025) and analyzed for sequence identity. The frequency of indicated lysine residues at these positions in PA is depicted. (B) H17N10 influenza virus vRNA-bound polymerase complex (PDB: 4WSB) showing PA (blue) residues 351, 353, 356, 357, 600, and 604 (orange red), which correspond to residues 356, 358, 361, 362, 605, and 609 in H1N1 PA, respectively. (C) HEK293T cells were transfected with influenza virus PB1, PB2, and NP expression plasmids, along with the pPol I-luc luciferase reporter plasmid, the pRL-TK Renilla luciferase internal control plasmid, and either WT PA, PA-C95A, PA-D108A, or a combination of PA-C95A and PA-D108A. At 24 h post-transfection, firefly and Renilla luciferase activities were measured using a dual-luciferase reporter assay system. Statistical analyses were performed using unpaired Student’s t-tests; *ns*, not significant; \*\**P*<0.01, \*\*\**P*<0.001.

**Figure S5.**
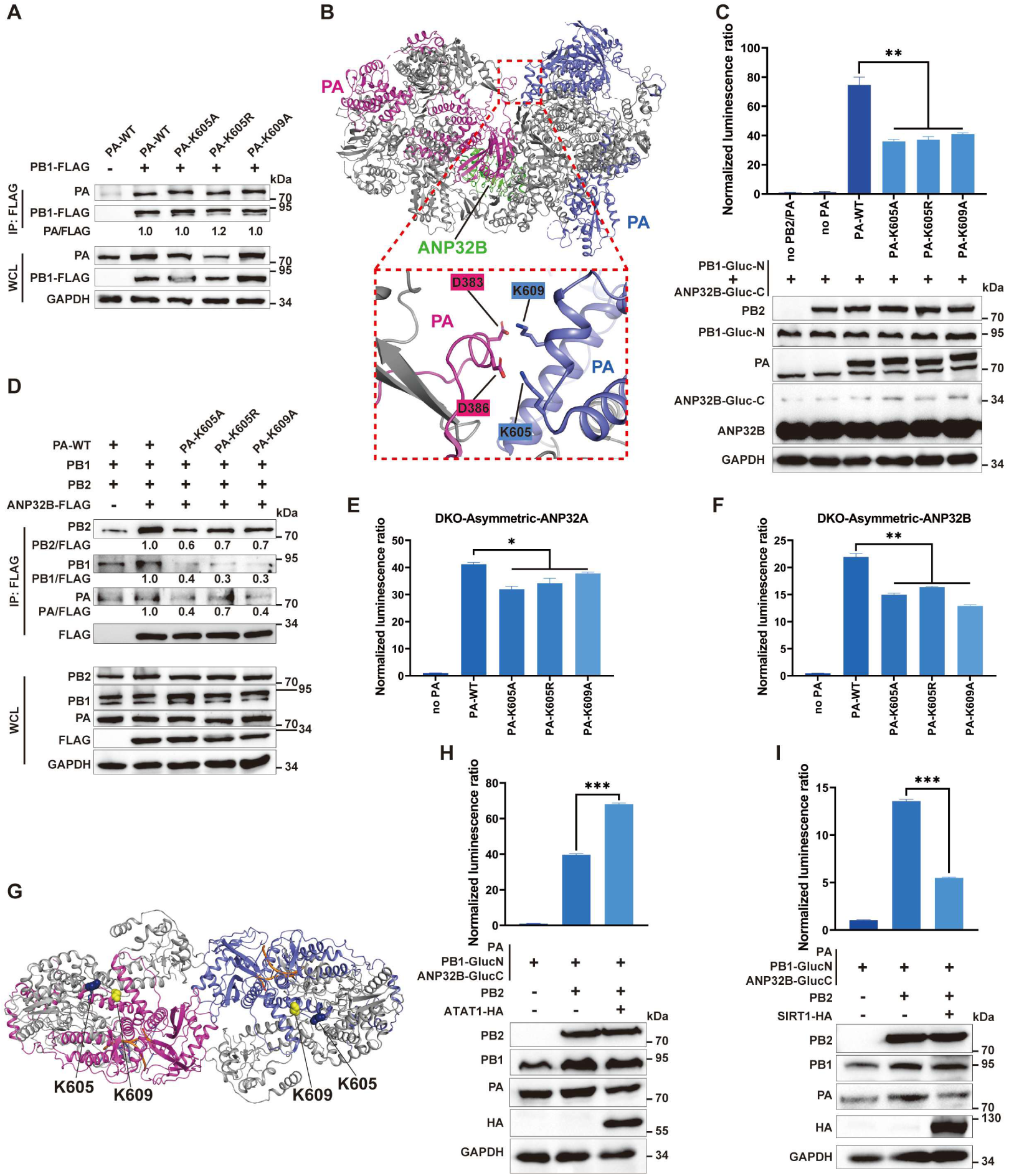
Lactylation of PA at K605/K609 facilitates ANP32B-mediated polymerase asymmetric dimerization. (A) HEK293T cells were co-transfected with FLAG-tagged PB1 and WT or PA-K605A/K605R/K609A mutants; anti-FLAG co-IP was performed at 24 h post-transfection, co-precipitated proteins were assessed by immunoblotting. Band intensities were quantified by ImageJ and normalized to that of WT-PA-transfected group. (B) H5N1 influenza virus polymerase asymmetric dimer (PDB: 8R1J) with ANP32B (green) showing PA K605 and K609. (C) HEK293T cells were co-transfected with PB2, PB1-Gluc-N, ANP32B-Gluc-C, and WT or PA-K605A/K605R/K609A mutants; Gaussia luciferase activity was measured at 24 h post-transfection; protein expression was detected by immunoblotting. (D) HEK293T cells were co-transfected with Flag-ANP32B, PB1, PB2, and WT or PA-K605A/K605R/K609A mutants; anti-Flag co-IP was performed at 24 h post-transfection; co-precipitated proteins were detected by immunoblotting. Band intensities were quantified by ImageJ and normalized to that of WT-PA-transfected group. (E-F) ANP32A/B double-knockout HEK293T cells were co-transfected with PB2, PB1-Gluc-N, ANP32A-Gluc-C (E) or ANP32B-Gluc-C (F), and WT or PA-K605A/K605R/K609A mutants; Gaussia luciferase activity was measured at 24 h post-transfection. (G) H3N2 influenza virus symmetric dimer (PDB: 6QX8) with PA K605 (blue) and K609 (yellow) depicted. (H-I) HEK293T cells were co-transfected with PB2, PB1-Gluc-N, ANP32B-Gluc-C, PA, and HA-ATAT1 (H) or HA-SIRT1 (I); Gaussia luciferase activity was measured at 24 h post-transfection; protein expression was detected by immunoblotting. Data are presented as mean ± SD and are representative of three independent experiments. Statistical analyses were performed using unpaired Student’s t-tests; *ns*, not significant; \**P*<0.05, \*\**P*<0.01, \*\*\**P*<0.001.

**Figure S6.**
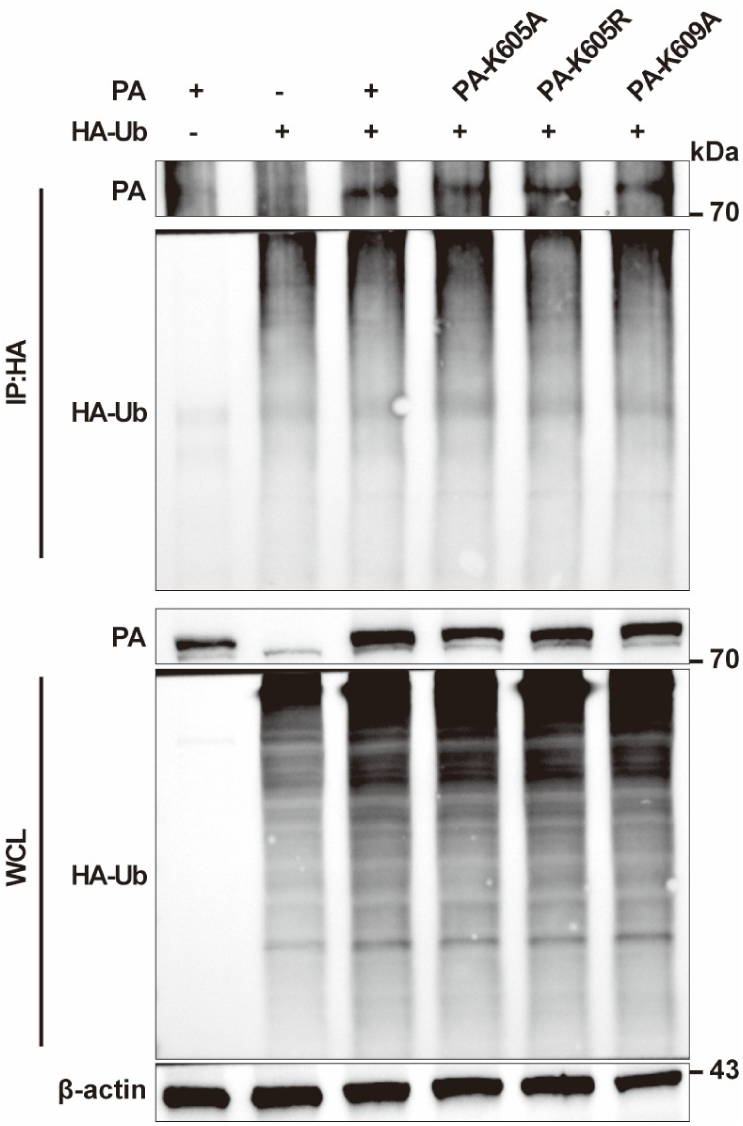
Mutations on K605/K609 residues of PA do not affect PA ubiquitination level. HEK293T cells were transfected with HA-tagged Ub and WT or PA-K605A/K605R/K609A mutants; anti-HA immunoprecipitation was performed at 24 h post-transfection.

## Notes

### Competing Interest Statement

The authors have declared no competing interest.

## References

1. Hutchinson EC, Amorim MJ, Yamauchi Y. 2025 Understanding Influenza. Methods Mol Biol. 2890:1–26.

2. Eisfeld AJ, Neumann G, Kawaoka Y. 2015 At the centre: influenza A virus ribonucleoproteins. Nat Rev Microbiol. 13(1):28–41.

3. Te Velthuis AJ, Fodor E. 2016 Influenza virus RNA polymerase: insights into the mechanisms of viral RNA synthesis. Nat Rev Microbiol. 14(8):479–93.

4. Deng T, Zhang L, Shi Y, Gao GF. 2025 In Transition: How Influenza Virus Switches from Transcription to Genome Replication. Annu Rev Virol. 12(1):239–58.

5. Nilsson-Payant BE, tenOever BR, Te Velthuis AJW. 2022 The Host Factor ANP32A Is Required for Influenza A Virus vRNA and cRNA Synthesis. J Virol. 96(4):e0209221.

6. Carrique L, Fan H, Walker AP, Keown JR, Sharps J, Staller E, Barclay WS, Fodor E, Grimes JM. 2020 Host ANP32A mediates the assembly of the influenza virus replicase. Nature. 587(7835):638–43.

7. Fan H, Walker AP, Carrique L, Keown JR, Serna Martin I, Karia D, Sharps J, Hengrung N, Pardon E, Steyaert J, Grimes JM, Fodor E. 2019 Structures of influenza A virus RNA polymerase offer insight into viral genome replication. Nature. 573(7773):287–90.

8. Dawson AR, Wilson GM, Coon JJ, Mehle A. 2020 Post-Translation Regulation of Influenza Virus Replication. Annu Rev Virol. 7(1):167–87.

9. Wei Y, Song J, Zhang J, Chen S, Yu Z, He L, Chen J. 2024 Exploring TRIM proteins’ role in antiviral defense against influenza A virus and respiratory coronaviruses. Front Cell Infect Microbiol. 14:1420854.

10. Husain M. 2024 Influenza A Virus and Acetylation: The Picture Is Becoming Clearer. Viruses. 16(1).

11. Dey S, Mondal A. 2024 Unveiling the role of host kinases at different steps of influenza A virus life cycle. J Virol. 98(1):e0119223.

12. Park ES, Dezhbord M, Lee AR, Kim KH. 2022 The Roles of Ubiquitination in Pathogenesis of Influenza Virus Infection. Int J Mol Sci. 23(9).

13. Hu J, Zhang L, Liu X. 2020 Role of Post-translational Modifications in Influenza A Virus Life Cycle and Host Innate Immune Response. Front Microbiol. 11:517461.

14. York IA, Stevens J, Alymova IV. 2019 Influenza virus N-linked glycosylation and innate immunity. Biosci Rep. 39(1).

15. Liang B, Fan M, Meng Q, Zhang Y, Jin J, Chen N, Lu Y, Jiang C, Zhang X, Zou Z, Ping J, Su J. 2024 Effects of the Glycosylation of the HA Protein of H9N2 Subtype Avian Influenza Virus on the Pathogenicity in Mice and Antigenicity. Transbound Emerg Dis. 2024:6641285.

16. Kim JI, Park MS. 2012 N-linked glycosylation in the hemagglutinin of influenza A viruses. Yonsei Med J. 53(5):886–93.

17. Su WC, Chen YC, Tseng CH, Hsu PW, Tung KF, Jeng KS, Lai MM. 2013 Pooled RNAi screen identifies ubiquitin ligase Itch as crucial for influenza A virus release from the endosome during virus entry. Proc Natl Acad Sci U S A. 110(43):17516–21.

18. Gao S, Wu J, Liu RY, Li J, Song L, Teng Y, Sheng C, Liu D, Yao C, Chen H, Jiang W, Chen S, Huang W. 2015 Interaction of NS2 with AIMP2 facilitates the switch from ubiquitination to SUMOylation of M1 in influenza A virus-infected cells. J Virol. 89(1):300–11.

19. Zhang D, Liang C, Wu C, Hawanga M, Wan S, Xu L, Zhang X, Liu Y, Hu F, Wang M, Wang X, Xu L, Huang X. 2025 Nonhistone lactylation: A hub for tumour metabolic reprogramming and epigenetic regulation. J Transl Med. 23(1):901.

20. Hu Y, He Z, Li Z, Wang Y, Wu N, Sun H, Zhou Z, Hu Q, Cong X. 2024 Lactylation: the novel histone modification influence on gene expression, protein function, and disease. Clin Epigenetics. 16(1):72.

21. Zhang D, Tang Z, Huang H, Zhou G, Cui C, Weng Y, Liu W, Kim S, Lee S, Perez-Neut M, Ding J, Czyz D, Hu R, Ye Z, He M, Zheng YG, Shuman HA, Dai L, Ren B, Roeder RG, Becker L, Zhao Y. 2019 Metabolic regulation of gene expression by histone lactylation. Nature. 574(7779):575–80.

22. Pang Y, Zhou Y, Wang Y, Fang L, Xiao S. 2024 Lactate-lactylation-HSPA6 axis promotes PRRSV replication by impairing IFN-β production. J Virol. 98(1):e0167023.

23. Zhang Y, Zhang X. 2024 Virus-Induced Histone Lactylation Promotes Virus Infection in Crustacean. Adv Sci (Weinh). 11(30):e2401017.

24. Yan Q, Zhou J, Gu Y, Huang W, Ruan M, Zhang H, Wang T, Wei P, Chen G, Li W, Lu C. 2024 Lactylation of NAT10 promotes N(4)-acetylcytidine modification on tRNA(Ser-CGA-1-1) to boost oncogenic DNA virus KSHV reactivation. Cell Death Differ. 31(10):1362–74.

25. Xin J, Wang C, Li Z, Gao W, Zhang W. 2025 Lactylation of the SARS-CoV-2 spike protein is required for viral infection. Signal Transduct Target Ther. 10(1):329.

26. Li Y, Wang Z, Wang J, Jiang Z, Chen M, Ai H, Ma C, Tong Q, Liu L, Velkov T, Sun H, Pu J, Liu J, Dai C, Sun Y. 2025 ARRDC4-mediated glycolysis enhances innate immunity to influenza A virus through fructose-1,6-bisphosphate. Proc Natl Acad Sci U S A. 122(35):e2512385122.

27. Zhang Y, Chang L, Xin X, Qiao Y, Qiao W, Ping J, Xia J, Su J. 2024 Influenza A virus-induced glycolysis facilitates virus replication by activating ROS/HIF-1α pathway. Free Radic Biol Med. 225:910–24.

28. Ren L, Zhang W, Zhang J, Zhang J, Zhang H, Zhu Y, Meng X, Yi Z, Wang R. 2021 Influenza A Virus (H1N1) Infection Induces Glycolysis to Facilitate Viral Replication. Virol Sin. 36(6):1532–42.

29. Thyrsted J, Storgaard J, Blay-Cadanet J, Heinz A, Thielke AL, Crotta S, de Paoli F, Olagnier D, Wack A, Hiller K, Hansen AL, Holm CK. 2021 Influenza A induces lactate formation to inhibit type I IFN in primary human airway epithelium. iScience. 24(11):103300.

30. Zhang W, Wang G, Xu ZG, Tu H, Hu F, Dai J, Chang Y, Chen Y, Lu Y, Zeng H, Cai Z, Han F, Xu C, Jin G, Sun L, Pan BS, Lai SW, Hsu CC, Xu J, Chen ZZ, Li HY, Seth P, Hu J, Zhang X, Li H, Lin HK. 2019 Lactate Is a Natural Suppressor of RLR Signaling by Targeting MAVS. Cell. 178(1):176–89.e15.

31. Kleinehr J, Bojarzyn CR, Schöfbänker M, Daniel K, Ludwig S, Hrincius ER. 2025 Metabolic interference impairs influenza A virus replication by dampening vRNA synthesis. Npj Viruses. 3(1):22.

32. Kleinehr J, Schöfbänker M, Daniel K, Günl F, Mohamed FF, Janowski J, Brunotte L, Boergeling Y, Liebmann M, Behrens M, Gerdemann A, Klotz L, Esselen M, Humpf HU, Ludwig S, Hrincius ER. 2023 Glycolytic interference blocks influenza A virus propagation by impairing viral polymerase-driven synthesis of genomic vRNA. PLoS Pathog. 19(7):e1010986.

33. Hoffmann E, Neumann G, Hobom G, Webster RG, Kawaoka Y. 2000 “Ambisense” approach for the generation of influenza A virus: vRNA and mRNA synthesis from one template. Virology. 267(2):310–7.

34. Tu S, Zou J, Xiong C, Dai C, Sun H, Luo D, Jin M, Chen H, Zhou H. 2024 Zinc-finger CCHC-type containing protein 8 promotes RNA virus replication by suppressing the type-I interferon responses. J Virol. 98(9):e0079624.

35. Kawakami E, Watanabe T, Fujii K, Goto H, Watanabe S, Noda T, Kawaoka Y. 2011 Strand-specific real-time RT-PCR for distinguishing influenza vRNA, cRNA, and mRNA. J Virol Methods. 173(1):1–6.

36. Sheppard CM, Goldhill DH, Swann OC, Staller E, Penn R, Platt OK, Sukhova K, Baillon L, Frise R, Peacock TP, Fodor E, Barclay WS. 2023 An Influenza A virus can evolve to use human ANP32E through altering polymerase dimerization. Nat Commun. 14(1):6135.

37. Mistry B, Long JS, Schreyer J, Staller E, Sanchez-David RY, Barclay WS. 2020 Elucidating the Interactions between Influenza Virus Polymerase and Host Factor ANP32A. J Virol. 94(3).

38. Cassonnet P, Rolloy C, Neveu G, Vidalain PO, Chantier T, Pellet J, Jones L, Muller M, Demeret C, Gaud G, Vuillier F, Lotteau V, Tangy F, Favre M, Jacob Y. 2011 Benchmarking a luciferase complementation assay for detecting protein complexes. Nat Methods. 8(12):990–2.

39. Li H, Liu C, Li R, Zhou L, Ran Y, Yang Q, Huang H, Lu H, Song H, Yang B, Ru H, Lin S, Zhang L. 2024 AARS1 and AARS2 sense L-lactate to regulate cGAS as global lysine lactyltransferases. Nature. 634(8036):1229–37.

40. Zong Z, Xie F, Wang S, Wu X, Zhang Z, Yang B, Zhou F. 2024 Alanyl-tRNA synthetase, AARS1, is a lactate sensor and lactyltransferase that lactylates p53 and contributes to tumorigenesis. Cell. 187(10):2375–92.e33.

41. Xie B, Zhang M, Li J, Cui J, Zhang P, Liu F, Wu Y, Deng W, Ma J, Li X, Pan B, Zhang B, Zhang H, Luo A, Xu Y, Li M, Pu Y. 2024 KAT8-catalyzed lactylation promotes eEF1A2-mediated protein synthesis and colorectal carcinogenesis. Proc Natl Acad Sci U S A. 121(8):e2314128121.

42. Stevaert A, Naesens L. 2016 The Influenza Virus Polymerase Complex: An Update on Its Structure, Functions, and Significance for Antiviral Drug Design. Med Res Rev. 36(6):1127–73.

43. Iuzzolino A, Pellegrini FR, Rotili D, Degrassi F, Trisciuoglio D. 2024 The α-tubulin acetyltransferase ATAT1: structure, cellular functions, and its emerging role in human diseases. Cell Mol Life Sci. 81(1):193.

44. Zhou Z, Yin X, Sun H, Lu J, Li Y, Fan Y, Lv P, Han M, Wu J, Li S, Liu Z, Zhao H, Sun H, Fan H, Wang S, Xin T. 2025 PTBP1 Lactylation Promotes Glioma Stem Cell Maintenance through PFKFB4-Driven Glycolysis. Cancer Res. 85(4):739–57.

45. Sun L, Zhang Y, Yang B, Sun S, Zhang P, Luo Z, Feng T, Cui Z, Zhu T, Li Y, Qiu Z, Fan G, Huang C. 2023 Lactylation of METTL16 promotes cuproptosis via m(6)A-modification on FDX1 mRNA in gastric cancer. Nat Commun. 14(1):6523.

46. Jin J, Bai L, Wang D, Ding W, Cao Z, Yan P, Li Y, Xi L, Wang Y, Zheng X, Wei H, Ding C, Wang Y. 2023 SIRT3-dependent delactylation of cyclin E2 prevents hepatocellular carcinoma growth. EMBO Rep. 24(5):e56052.

47. Moreno-Yruela C, Zhang D, Wei W, Bæk M, Liu W, Gao J, Danková D, Nielsen AL, Bolding JE, Yang L, Jameson ST, Wong J, Olsen CA, Zhao Y. 2022 Class I histone deacetylases (HDAC1-3) are histone lysine delactylases. Sci Adv. 8(3):eabi6696.

48. Davenport AM, Huber FM, Hoelz A. 2014 Structural and functional analysis of human SIRT1. J Mol Biol. 426(3):526–41.

49. Xu K, Zhang K, Wang Y, Gu Y. 2024 Comprehensive review of histone lactylation: Structure, function, and therapeutic targets. Biochem Pharmacol. 225:116331.

50. Nilsson-Payant BE, Sharps J, Hengrung N, Fodor E. 2018 The Surface-Exposed PA(51-72)-Loop of the Influenza A Virus Polymerase Is Required for Viral Genome Replication. J Virol. 92(16).

51. Deng T, Sharps J, Fodor E, Brownlee GG. 2005 In vitro assembly of PB2 with a PB1-PA dimer supports a new model of assembly of influenza A virus polymerase subunits into a functional trimeric complex. J Virol. 79(13):8669–74.

52. Chen D, Zhao G, Zhou J, Sun P, He S, Lv C, Chen Y, Zhu S, Gao M, Guo G. 2026 Delactylation of viral proteins by SIRT1 suppresses influenza A virus replication. mBio.e0248925.

53. Wan N, Wang N, Yu S, Zhang H, Tang S, Wang D, Lu W, Li H, Delafield DG, Kong Y, Wang X, Shao C, Lv L, Wang G, Tan R, Wang N, Hao H, Ye H. 2022 Cyclic immonium ion of lactyllysine reveals widespread lactylation in the human proteome. Nat Methods. 19(7):854–64.

54. Giese S, Ciminski K, Bolte H, Moreira É A, Lakdawala S, Hu Z, David Q, Kolesnikova L, Götz V, Zhao Y, Dengjel J, Chin YE, Xu K, Schwemmle M. 2017 Role of influenza A virus NP acetylation on viral growth and replication. Nat Commun. 8(1):1259.

55. Hu J, Liu X. 2015 Crucial role of PA in virus life cycle and host adaptation of influenza A virus. Med Microbiol Immunol. 204(2):137–49.

56. Hatakeyama D, Shoji M, Ogata S, Masuda T, Nakano M, Komatsu T, Saitoh A, Makiyama K, Tsuneishi H, Miyatake A, Takahira M, Nishikawa E, Ohkubo A, Noda T, Kawaoka Y, Ohtsuki S, Kuzuhara T. 2022 Acetylation of the influenza A virus polymerase subunit PA in the N-terminal domain positively regulates its endonuclease activity. Febs j. 289(1):231–45.

57. Chen H, Qian Y, Chen X, Ruan Z, Ye Y, Chen H, Babiuk LA, Jung YS, Dai J. 2019 HDAC6 Restricts Influenza A Virus by Deacetylation of the RNA Polymerase PA Subunit. J Virol. 93(4).

58. Zhang M, Zeng Z, Chen X, Wang G, Cai X, Hu Z, Gu M, Hu S, Liu X, Wang X, Peng D, Hu J, Liu X. 2025 Phosphorylation of PA at serine 225 enhances viral fitness of the highly pathogenic H5N1 avian influenza virus in mice. Vet Microbiol. 302:110400.

59. Liu L, Madhugiri R, Saul VV, Bacher S, Kracht M, Pleschka S, Schmitz ML. 2023 Phosphorylation of the PA subunit of influenza polymerase at Y393 prevents binding of the 5’-termini of RNA and polymerase function. Sci Rep. 13(1):7042.

60. Wang ZA, Cole PA. 2020 The Chemical Biology of Reversible Lysine Post-translational Modifications. Cell Chem Biol. 27(8):953–69.

61. Vu LD, Gevaert K, De Smet I. 2018 Protein Language: Post-Translational Modifications Talking to Each Other. Trends Plant Sci. 23(12):1068–80.

62. Günl F, Krischuns T, Schreiber JA, Henschel L, Wahrenburg M, Drexler HCA, Leidel SA, Cojocaru V, Seebohm G, Mellmann A, Schwemmle M, Ludwig S, Brunotte L. 2023 The ubiquitination landscape of the influenza A virus polymerase. Nat Commun. 14(1):787.

63. Even A, Morelli G, Broix L, Scaramuzzino C, Turchetto S, Gladwyn-Ng I, Le Bail R, Shilian M, Freeman S, Magiera MM, Jijumon AS, Krusy N, Malgrange B, Brone B, Dietrich P, Dragatsis I, Janke C, Saudou F, Weil M, Nguyen L. 2019 ATAT1-enriched vesicles promote microtubule acetylation via axonal transport. Sci Adv. 5(12):eaax2705.

64. Glon D, Vilmen G, Perdiz D, Hernandez E, Beauclair G, Quignon F, Berlioz-Torrent C, Maréchal V, Poüs C, Lussignol M, Esclatine A. 2022 Essential role of hyperacetylated microtubules in innate immunity escape orchestrated by the EBV-encoded BHRF1 protein. PLoS Pathog. 18(3):e1010371.

65. Qin Z, Fang X, Sun W, Ma Z, Dai T, Wang S, Zong Z, Huang H, Ru H, Lu H, Yang B, Lin S, Zhou F, Zhang L. 2022 Deactylation by SIRT1 enables liquid-liquid phase separation of IRF3/IRF7 in innate antiviral immunity. Nat Immunol. 23(8):1193–207.

66. Tang HM, Gao WW, Chan CP, Cheng Y, Deng JJ, Yuen KS, Iha H, Jin DY. 2015 SIRT1 Suppresses Human T-Cell Leukemia Virus Type 1 Transcription. J Virol. 89(16):8623–31.

67. Zhang Y, Li J, Qiu Z, Huang L, Yang S, Li J, Li K, Liang Y, Liu X, Chen Z, Li J, Zhou B. 2024 Insights into the mechanism of action of pterostilbene against influenza A virus-induced acute lung injury. Phytomedicine. 129:155534.

68. Simeonova L, Leseva M, Stoyanov K, Mai A, Saso L, Dimitrova P. 2025 Neutrophils shape the therapeutic efficacy of sirtuin 1 activity modulators in murine influenza virus infection. Biochim Biophys Acta Mol Basis Dis. 1871(7):167915.

69. Staller E, Carrique L, Swann OC, Fan H, Keown JR, Sheppard CM, Barclay WS, Grimes JM, Fodor E. 2024 Structures of H5N1 influenza polymerase with ANP32B reveal mechanisms of genome replication and host adaptation. Nat Commun. 15(1):4123.

70. Arragain B, Krischuns T, Pelosse M, Drncova P, Blackledge M, Naffakh N, Cusack S. 2024 Structures of influenza A and B replication complexes give insight into avian to human host adaptation and reveal a role of ANP32 as an electrostatic chaperone for the apo-polymerase. Nat Commun. 15(1):6910.

